# DNA methylation in human gastric epithelial cells allows cell type-related plasticity and defines regional identity

**DOI:** 10.1101/2022.03.29.486203

**Authors:** Kristin Fritsche, Francesco Boccellato, Philipp Schlaermann, Max Koeppel, Christian Denecke, Ivo Gut, Thomas F. Meyer, Hilmar Berger

**Affiliations:** Department of Molecular Biology, Max Planck Institute for Infection Biology, 10117 Berlin, Germany; Ludwig Institute for Cancer Research, Nuffield Department of Clinical Medicine, University of Oxford, United Kingdom.; Center for Bariatric and Metabolic Surgery, Center of Innovative Surgery (ZIC), Department of Surgery, Campus Virchow Klinikum and Campus Mitte, Charité-Universitätsmedizin Berlin, Berlin, Germany.; Centro Nacional de Análisis Genómico (CNAG-CRG), Barcelona, Spain; Laboratory of Infection Oncology, Institute of Clinical Molecular Biology, Christian Albrecht University of Kiel and University Hospital Schleswig-Holstein – Campus Kiel, Germany

**Author notes:** Corresponding authors: Dr. Hilmar Berger Institute of Clinical Molecular Biology University Hospital Schleswig-Holstein – Campus Kiel Rosalind-Franklin-Straße 12 24105 Kiel Germany Prof. Dr. Thomas F. Meyer Max Planck Institute for Infection Biology Charitéplatz 1 10117 Berlin Germany.

**Keywords:** epigenetic regulation, human stomach, gastric epithelial development, gastric cancer, intestinal metaplasia

## Abstract

**Background:** Epigenetic modifications in mammalian DNA are commonly manifested by DNA methylation. In the stomach, altered DNA methylation patterns have been observed following chronic *Helicobacter pylori* infections and in gastric cancer. In the context of epigenetic regulation, the regional nature of the stomach has been rarely considered in detail.

**Results:** Here, we describe the DNA methylation landscape across the phenotypically different regions of the healthy human stomach (i.e., antrum, corpus, fundus) together with the corresponding transcriptomes. We show that stable regional DNA methylation differences translate to a limited extent into regulation of the transcriptomic phenotype, indicating a largely permissive epigenetic regulation. We identify a small number of transcription factors with novel region-specific activity and likely epigenetic impact in the stomach, including GATA4, IRX5, IRX2, PDX1, and CDX2. Detailed analysis of the Wnt pathway reveals differential regulation along the craniocaudal axis, which involves non-canonical Wnt signaling in determining cell fate in the proximal stomach.

**Conclusions:** By extending our analysis to pre-neoplastic lesions and gastric cancers, we conclude that epigenetic dysregulation already characterizes intestinal metaplasia as a founding basis for functional changes in gastric cancer. Finally, our study provides a well-defined resource of regional stomach transcription and epigenetics as a starting point for further studies.

## Background

Methylation of cytosines in the DNA represents the most studied heritable epigenetic modification that is highly important in embryonic development, tissue formation and cellular state maintenance (Greenberg and Bourc’his, 2019). DNA methylation patterns are established early in development after two waves of erasure in gametogenesis and early embryogenesis (Monk, 2015). Methylation of promoter and enhancer regions during development often leads to stable repression of transcription factors (TFs) (Fouse et al., 2008; Ng et al., 2008) and is essential to stabilize cell lineages and phenotypes (Bird, 2002; Bock et al., 2012). Once established, these patterns remain stable throughout life. They can, however, undergo stochastic or deterministic changes, which mostly depend on the epigenetic context of a given site (Beerman et al., 2013; Marttila et al., 2015; Schlesinger et al., 2007; Zhou et al., 2018).

The human stomach consists of distinct regions along its cranial-caudal axis (gastric cardia, fundus, corpus or body and antrum), characterized by differences in mucosa phenotype, cellular composition and function (Hoffmann, 2015). These characteristics arise during development and are stable over time (Kim and Shivdasani, 2016). The role of DNA methylation in the establishment and homeostasis of these regional differences has not been elucidated in detail so far, despite some preliminary regional observations (Huang et al., 2018). Aberrant DNA methylation has been linked to several pathological conditions, including cancer (Cancer Genome Atlas Research Network, 2014; Yoda et al., 2015). In particular, it is also a hallmark of gastric adenocarcinoma that arises from a transformed healthy glandular epithelium (Yoda et al., 2015). Gastric cancer (GC) development is predominantly associated with chronic *Helicobacter pylori* infection (Uemura et al., 2001). Notably, changes in DNA methylation are already observed early in infected tissues and premalignant lesions, including intestinal metaplasia (IM) (Huang et al., 2012, 2018; Niwa et al., 2010). Although our understanding of DNA methylation dynamics increased drastically in the past years along with improved methods of its detection (Bibikova et al., 2011; Moran et al., 2016), the mechanisms leading to aberrant DNA methylation in GC, apart from tumors with gastric CpG island methylator phenotype (CIMP) (Hughes et al., 2013), remain largely unknown. Recent advances in primary cell culture techniques allow to maintain and expand pure epithelial cells derived from healthy or diseased tissue (Boccellato et al., 2018; Nanki et al., 2018). Primary human gastric epithelial cells derived from the antrum and corpus can be cultivated in the presence of Wnt Family Member 3A (WNT3A) and R-Spondin 1 (RSPO1), where they maintain region-specific transcriptional programs (Schlaermann et al., 2016). Further, their differentiation to pit cells can be induced upon the withdrawal of the WNT3A and RSPO1 morphogens (Boccellato et al., 2018). To understand the characteristics and dynamics of DNA methylation in human gastric epithelial cells, we cultivated primary cells, which were obtained from healthy antrum, corpus, and fundus, as plane mucosoids in the presence and absence of WNT3A and RSPO1 (Boccellato et al., 2018), followed by analyses of DNA methylation and gene expression patterns. We ran detailed bioinformatic analyses of epigenetic and transcriptional differences, comparing different stomach regions, differentiation states, and pathological states. Our data were integrated with published methylomes of healthy gastrointestinal tissues, novel *ex vivo*-cultivated gastric primary cells of IM, and public data sets of IM tissue and GC. We describe an epigenetic landscape that allows differentiation plasticity with only minor restrictions underlying the regional phenotypic differences in the stomach.

## Results

### *In-vitro* model of primary human gastric epithelial cells recapitulates regional DNA methylation profiles

To investigate cell-type-specific DNA methylation patterns, we isolated human gastric epithelial cells of the antrum, corpus and fundus from sleeve resections and cultivated them as mucosoids (Boccellato et al., 2018). Stem cell-enriched cell populations of the antrum, corpus, and fundus were maintained in a medium containing WNT3A and RSPO1 (+W/R) or differentiated *in vitro* to pit cells after removal of both factors (-W/R, Fig. 1A). Genome-wide DNA methylation analysis of three biological replicates revealed that cells from different regions preserved a clear regional identity (Fig. 1B-C), while clusters distinguishing -W/R from +W/R cells were not observed. We sought to identify CpGs showing regional methylation differences irrespective of +W/R and -W/R treatment. Out of 738,115 CpGs, 3,703 CpGs were found to be differentially methylated (DM) by comparing each two stomach regions (i.e., inter-regional comparison; delta beta > 0.2 or < −0.2, FDR < 5 %) (Fig. 1C, Additional Fig. 1A, Additional Table 1). Inter-regional methylation differences determined *in vitro* correlated very well with those determined in *in vivo* samples from similar regions (Huang et al., 2018) (Fig. 1D, Additional Fig. 1B). Thus, the *in vitro* mucosoids maintain their region-specific DNA methylation profile. Because DNA methylation changes upon differentiation are known to be less prominent (Sheaffer et al., 2014), we performed a sensitivity analysis lowering the thresholds of delta beta (i.e., the difference in DM) and p-value in the comparisons between -W/R and +W/R conditions for each stomach region (Additional Fig. 1C). In agreement with previous findings, no stem cell- or differentiation-specific gene was found to be consistently affected by DM across regions (Additional Fig. 1D). Instead, we detected only single CpGs with weak differences below the confidence threshold. It should be mentioned, though, that in the corpus, the suggested stem cell marker *TNFRSF19* (TNF Receptor Superfamily Member 19, also known as TROY) was hypermethylated in a fraction of differentiated cells compared to undifferentiated cells.

**Figure 1:**
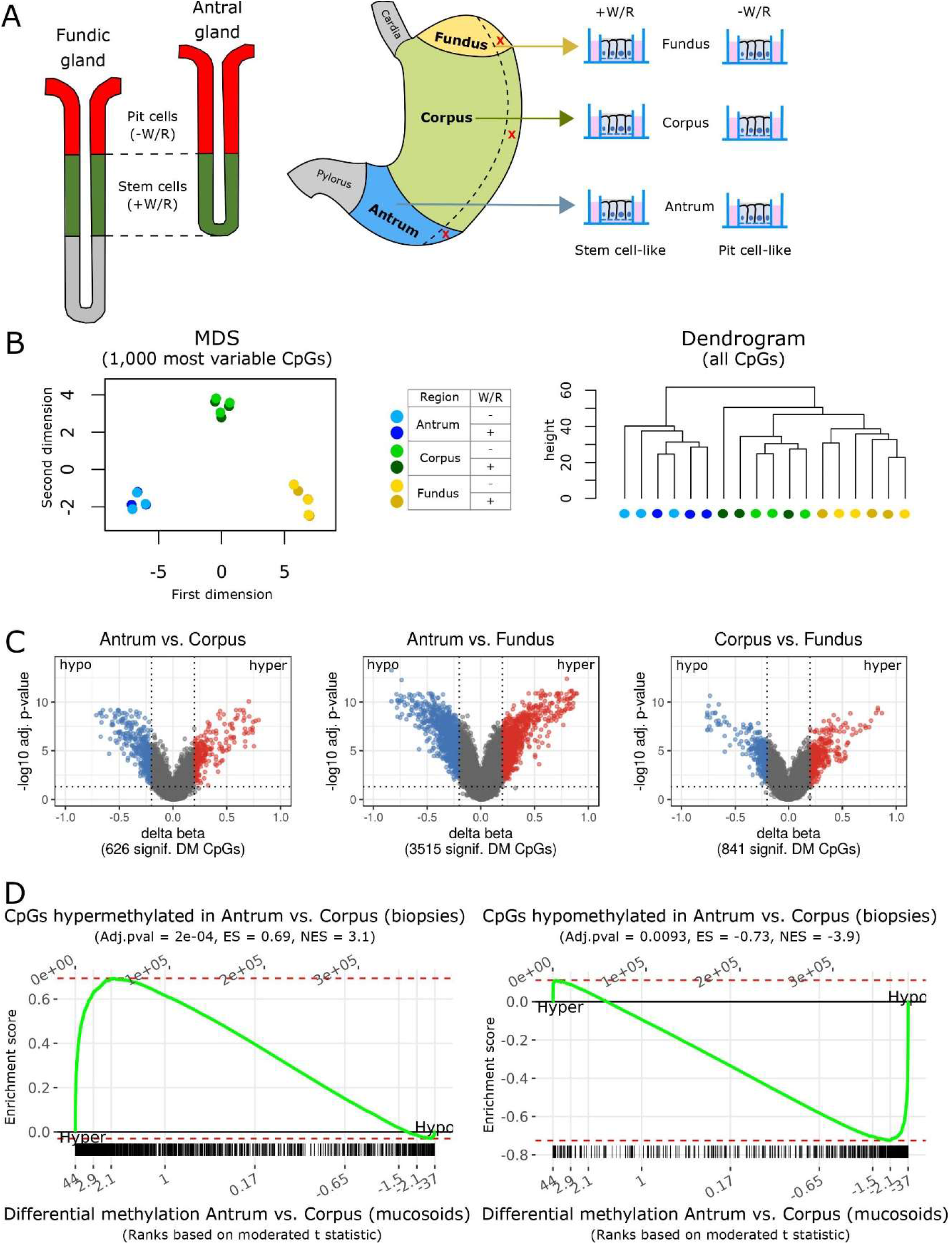
Global patterns of DNA methylation in the stomach. **A)** Left: schematic representation of the fundic and the antral gland types and the respective location of pit cells and stem cells. Right: an experimental overview. Following sleeve resection, sample tissues (red X) from the antrum, corpus, and fundus were cultivated as mucosoids using air-liquid-interface cell culture inserts. The removal or presence of WNT3A and RSPO1 (W/R) from the cell culture medium enriched for differentiated pit-like cells (-W/R) or stem cell-like cell populations (+W/R), respectively. **B)** Left: multidimensional scaling (MDS) of methylation proportions (beta) based on the 1,000 most variable CpGs in the normalized data set; samples from three biological replicates under the indicated conditions. Right: hierarchical cluster analysis (dendrogram) of all CpGs (n = 738,115) in the normalized data set. **C)** Volcano plots representing stomach inter-regional comparisons (+W/R and -W/R combined). The dashed horizontal and vertical lines indicate cutoffs of FDR < 0.05 and delta beta between −0.2 and −0.2, respectively. Red dots refer to hypermethylated CpGs (hyper) and blue dots to hypomethylated CpGs (hypo). **D)** Enrichment of sets of CpGs hypermethylated (left) or hypomethylated (right) in antrum-compared to corpus biopsies relative to differential methylation of CpGs between antrum- and corpus mucosoids. Moderated t-scores were used for DM CpG ranking. ES - enrichment score; NES - normalized enrichment score. Hyper, Hypo indicate the direction of differential methalytion in the antrum vs. corpus mucosoids comparison.

### Regional DNA methylation differences in the gastric epithelium suggest stem cell plasticity

In total, 170 and 159 genes were found to be affected by hypermethylation or hypomethylation, respectively, comparing any two stomach regions (Additional Fig 2A, Additional Table 2). The top 15 most affected genes between all regions, ordered by their percentage of DM CpGs out of all interrogated CpGs per gene, include several well-known developmental (homeodomain-) TFs (Fig. 2A). A gene set enrichment analysis of all DM genes revealed that almost half (147 out of 315 unique genes) of these genes are indeed involved in developmental processes like tissue development and pattern specification (Fig. 2B, Additional Table 3). Strikingly, only 11 genes of all DM genes demonstrated gene expression differences (Additional Fig. 2B). Of these, only four genes were found where promoter methylation negatively or positively correlated with gene expression (Fig. 2C). Interestingly, among genes most affected by differential methylation, we identified the pan-stomach developmental regulator GATA Binding Protein 4 (*GATA4*), Pancreatic And Duodenal Homeobox 1 (*PDX1*), which is a transcription factor essential for the development of the pancreas and gastroduodenal junction and expressed in gastric antrum (Holland et al., 2013; Offield et al., 1996; Stoffers et al., 1999), as well as Iroquois Homeobox 2 and 5 (*IRX2* and *IRX5*), which have been implicated in the specification of stomach fundus in mice (McCracken et al., 2017) (Fig. 2A). Furthermore, we detected the following inter-regionally DM genes: the Meis Homeobox family proteins *MEIS1* and *MEIS2*, which cooperatively bind DNA with several other homeodomain-containing TFs, like PDX1 or GATA TFs (Schulte and Geerts, 2019), the intestinal master regulator Caudal Type Homeobox 2 (*CDX2*) (Stringer et al., 2012), and the transcription factor SIM BHLH Transcription Factor 2 (*SIM2*). Using additional *in vivo* and *in vitro* data sets (see Methods, Additional table 7), we validated differential methylation of these genes at their respective promoter and enhancer sites (Additional Table 4) and evaluated their gene expression (Fig. 2D, Additional Fig. 3). Our analysis revealed that all of these DM sites bind Enhancer Of Zeste 2 Polycomb Repressive Complex 2 Subunit (EZH2), and most of them also SUZ12 Polycomb Repressive Complex 2 Subunit (SUZ12). EZH2 is the functional enzymatic subunit of the polycomb repressive complex 2 (PRC2), which catalyzes the methylation of lysine 27 of histone 3 (H3K27), leading to transcriptional repression (Laugesen et al., 2019). PRC2 proteins, such as EZH2 and SUZ12, play a critical role in stem cell maintenance, lineage specification, and differentiation (Batool et al., 2019; Rajasekhar and Begemann, 2007). We performed locus overlap analysis (Sheffield and Bock, 2016) to identify known TF binding sites (TFBS) across discovered stomach-associated DM CpGs corresponding to published embryonic stem cell data sets (Roadmap Epigenomics Consortium et al., 2015; Siggens and Ekwall, 2014). In line with the enrichment at binding sites for EZH2 and SUZ12 (Fig. 2E), we also detected a regional enrichment of DM CpGs at sites marked with trimethylation at lysine 9 and 27 of histone 3 (H3K9me3 and H3K27me3). In contrast, these genomic features were not enriched by comparing DM CpGs between the stomach and adjacent tissues (Fig. 2E). Instead, H3K4me1, a histone modification characteristic of enhancers (Heintzman et al., 2007; Pekowska et al., 2011), was enriched. In contrast to the inter-regional differences, we further detected that differences with other tissues preferentially occur at open sea CpGs (Additional Fig. 2C), which might be related to cell-type-specific differences in A-B compartments (Fortin and Hansen, 2015; Lieberman-Aiden et al., 2009). Many stomach-specific genes are expressed in individual cell lineages, predominantly in selected areas of the stomach, for example, parietal cells in the corpus and fundus and G-cells in the antrum. We determined the percentage of DM among stomach-specific genes (Fig. 2F) and found that they are rarely affected inter-regionally in the stomach in contrast to the comparisons between different tissues. Together, the small number of stomach cell lineage-specific genes affected by differential methylation and the small number of regulated TFs indicate that the regulation of transcription programs within the stomach is rather permissive and does not include stringent specification of lineages.

**Figure 2:**
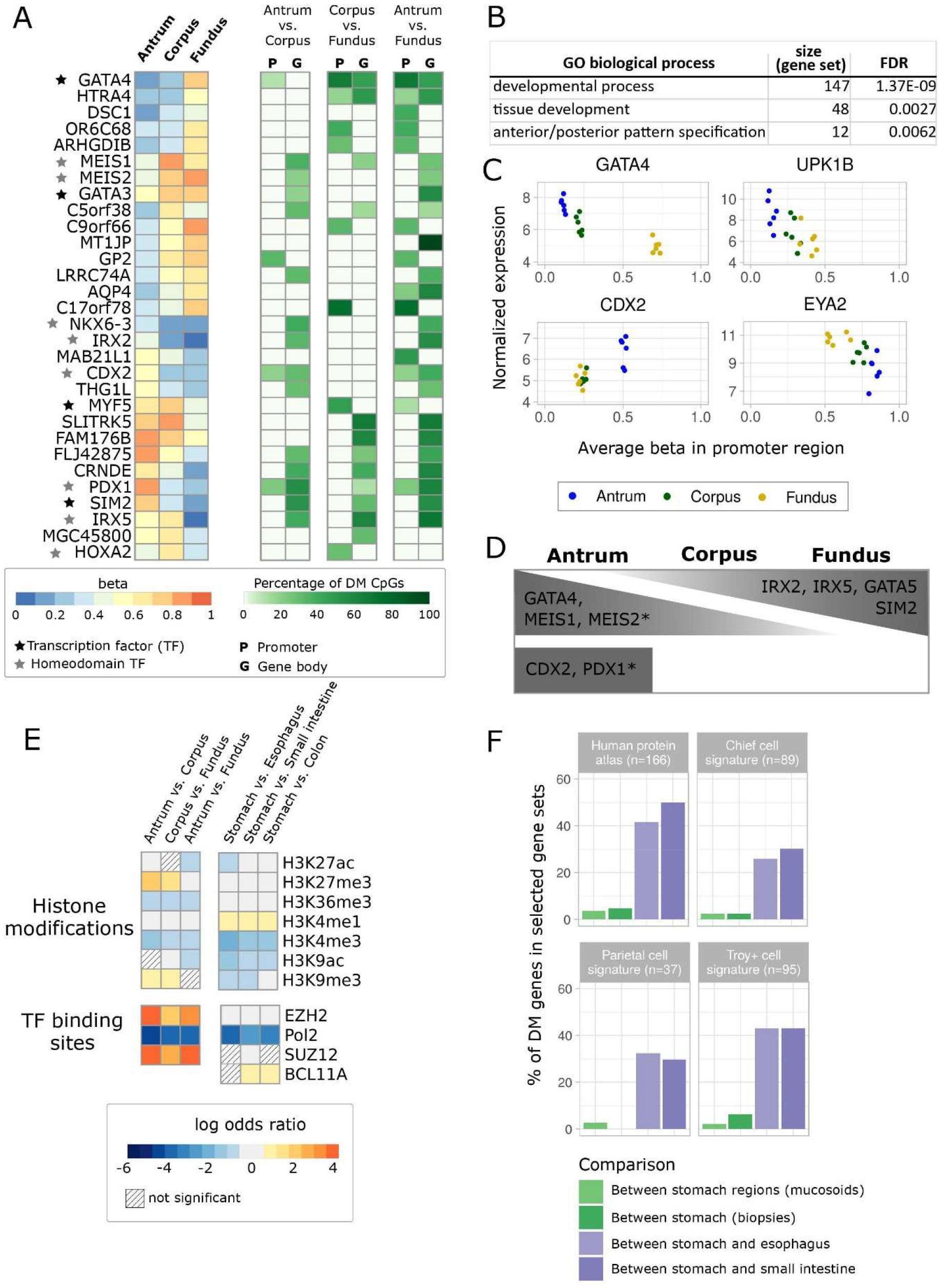
Differential methylation within the stomach and between the stomach and adjacent tissues. **A)** Displayed are the top 15 DM genes in all comparisons, ranked by the highest number of DM CpGs normalized to total CpGs/gene. Left: a heatmap of average DNA methylation (beta) values per gene. Color code ranges from 0 (unmethylated, blue) to 1 (methylated, red). Stars refer to (homeodomain) transcription factors (TFs). Right: the percentage of significantly DM CpGs out of the total number of interrogated CpGs per gene; color code ranges from white to dark green refers to percentage. **B)** Table of selected gene ontology (GO) terms enriched in antrum vs. fundus, determined with StringDB (Szklarczyk et al., 2015) **C)** Average promoter methylation is plotted against the normalized gene expression values. The selected genes showed negative (GATA4,UPK1B,EYA2) and positive (CDX2) correlations between promoter methylation and gene expression. **D)** Schematic representation of expression patterns of several DM TFs in different stomach regions, based on mucosoid and biopsy data sets (Table 3, Additional Fig. 3). Asterisks indicate genes not differentially expressed in the mucosoid data set **E)** Enrichment and depletion of DM CpGs according to different genomic features – heatmap comparisons between stomach regions and between the stomach and adjacent tissues. Shown are significantly enriched (log odds ratio > 0.6) and depleted (log odds ratio < −0.6) histone modifications and TF binding sites (FDR < 0.05). Color code ranges from blue (depleted) to red (enriched); light grey refers to a log odds ratio between −0.6 and 0.6 ; shaded squares display non-significant results; Pol2 - Polymerase 2 subunit. **F)** Stomach- and stomach cell type-specific gene sets - percentages of DM genes between stomach regions (green) and between the stomach and adjacent tissues (purple).

### Small but specific gene expression differences define the regional identity of stem cells in-vitro

In contrast to methylation levels, the global transcriptomic landscape of +W/R and -W/R stomach regions appeared to be governed by changes between stem cells and pit cell-like cell populations (Fig. 3A and 3B, top). However, we also identified regional gene expression differences between the antrum and the corpus *in vitro,* which correlated well with those *in vivo* (Additional Fig. 4A). Of note, the expression of marker genes of dominant chief- and parietal cells in the corpus and fundus was not induced *in vitro* (Additional Fig. 4B). The presence of antral endocrine G and D cells was confirmed by *GAST* (Gastrin) and *SST* (Somatostatin) expression, respectively (Additional Fig. 4B). Most proposed stem cell markers in the stomach, including the antral marker Leucine-Rich Repeat Containing G Protein-Coupled Receptor 5 (*LGR5*), the corpus markers *TNFRSF19* (*TROY*), SRY-Box Transcription Factor 2 (*SOX2*), and RUNX Family Transcription Factor 1 (*RUNX1*), were expressed as reported (Arnold et al., 2011; Barker et al., 2010; Matsuo et al., 2017; Stange et al., 2013) in the +W/R cultures (Fig. 3C). Despite these differences in stem cell gene expression, only a few other genes were differentially expressed inter-regionally (Fig. 3B, bottom). The antrum vs. fundus comparison revealed the relatively largest number of differences, including genes that were also DM (Fig. 3D, Additional Fig. 2B). A gene set enrichment analysis of the fundus compared to antrum further indicated a higher expression of genes associated with complement activation and adaptive and innate immune responses (Fig. 3E, Additional Table 5). Interestingly, three of the leading-edge genes in adaptive and innate immune response gene sets, Beta-2-Microglobulin (*B2M*), Protein Tyrosine Kinase 2 Beta (*PTK2B*), and BCL6 Transcription Repressor (*BCL6*), were also hypermethylated in the antrum compared to fundus (Additional Table 1). Hypermethylation at *B2M* affected the gene body with annotated Fantom5 enhancer site (Lizio et al., 2015), whereas *PTK2B* and *BCL6* hypermethylation occurred in the promoter region. Together, transcriptome analysis of stomach cells under +W/R and –W/R conditions *in vitro* present broadly similar profiles with only minor transcriptional differences.

**Figure 3:**
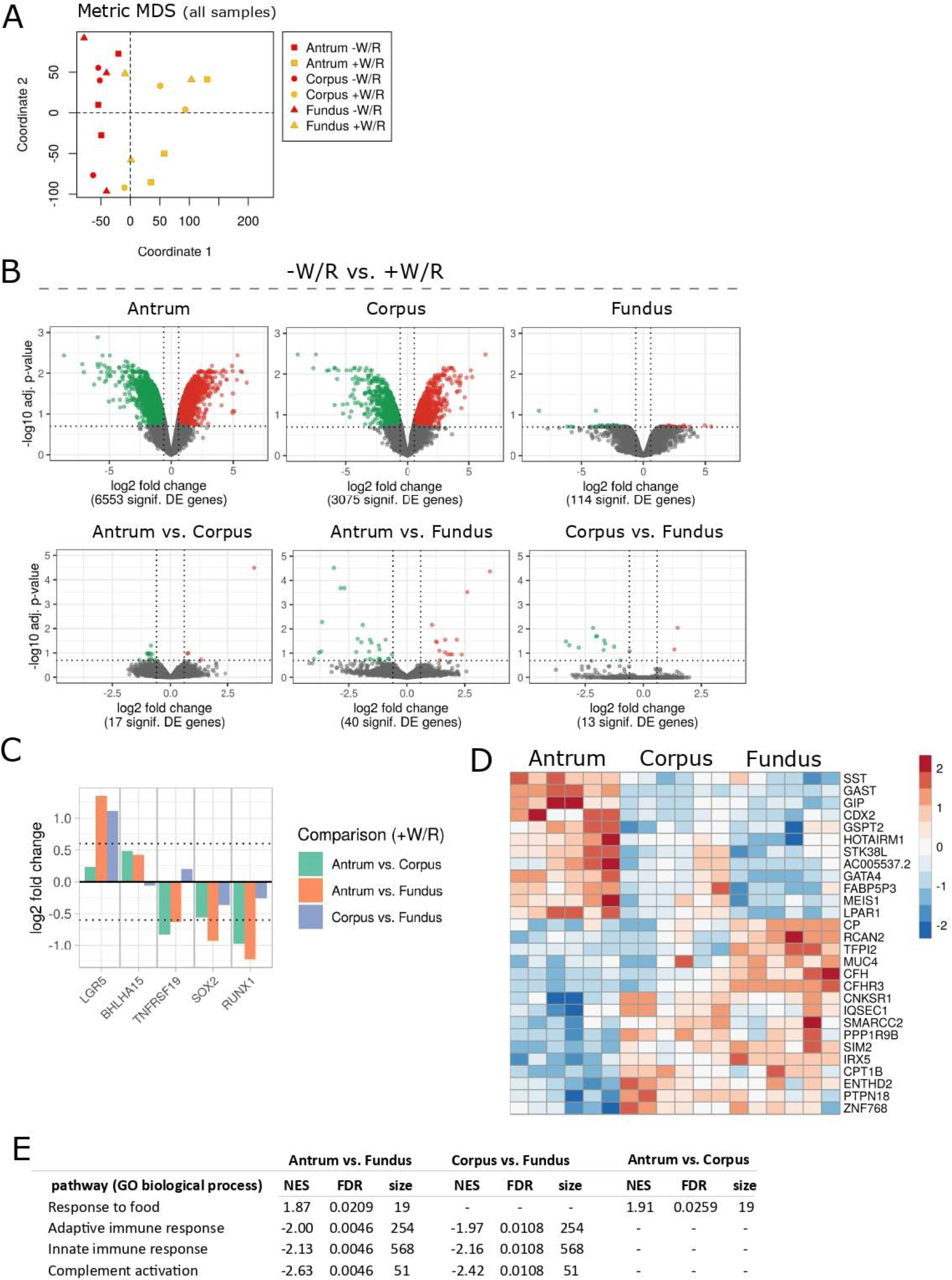
Differential gene expression in the stomach. **A**) MDS of gene expression values in all samples. **B)** Volcano plots of gene expression log2 fold changes between -W/R and +W/R (top) and between the stomach regions (+W/R and -W/R combined per region) (bottom). The dashed horizontal and vertical lines indicate cutoffs of adjusted p-value (FDR) < 0.2 and a log2 fold-change at −0.6 and 0.6, respectively. **C)** Regional comparison of differentially regulated stem cell genes in undifferentiated samples (+W/R), (p < 0.05). **D)** Heatmap of the top 15 up- and down-regulated genes (FDR < 0.2) in all regional comparisons. The normalized expression is represented by row-wise standardization (z-score). Color code ranges from blue (low expression) to red (high expression) **E)** Selected GO terms of biological process based on the GSEA of antrum vs. fundus. NES - normalized enrichment score.

### Wnt responsiveness decreases from the antrum to the fundus

The withdrawal of the morphogens W/R induced differentiation to pit cells in all stomach regions (Fig. 4A). Differentiated pit cells characteristically expressed Gastrokine 1 and 2 (*GKN1* and *GKN2*) and Mucin 5AC (*MUC5AC*), while the stem cell-enriched population characteristically expressed Mucin 6 (*MUC6*) and *CD44*. Although canonical Wnt signaling was active in all three stomach regions (Fig. 4B), we detected regional differences in the expression of Wnt pathway genes, with antrum showing the highest number of up and downregulated genes followed by corpus and fundus, and only a small number of differentially expressed (DE) genes shared by all three regions (Additional Fig. 4C). Thus, we determined the expression levels of crucial Wnt signaling genes upon induced differentiation. While the major Wnt target Axin 2 (*AXIN2*) was equally downregulated in differentiating mucosoids, indicating active Wnt signaling in cells from all regions, other genes demonstrated a region-dependent effect (Additional Fig. 4C-D). The differentiation effect on the expression of LGR5 and Transcription Factor 7 Like 1 (*TCF7L1*, one of the four TCF/LEF proteins that mediate Wnt signaling) weakened along the inferior-superior axis, while the remaining members of the family (TCF7, TCF7L2 and LEF1) displayed only minor or no change along this axis (Fig. 4C, Additional Fig. 4C-D). Interestingly, we detected an opposite trend in the expression of the non-canonical Wnt signaling genes Wnt Family Member 5A (*WNT5A*) and its putative receptor Receptor Tyrosine Kinase Like Orphan Receptor 2 (*ROR2*) (Oishi et al., 2003; Saldanha et al., 1998), whose expression in the corpus and fundus was higher than the antrum, regardless of the differentiation state (Fig. 4C). Moreover, we detected DM of *WNT5A*, *TCF7L1*, and *TCF7,* comparing the antrum and the fundus (Fig. 4D). In this comparison, *WNT5A* and *TCF7* were hypermethylated and hypomethylated, respectively, at one CpG, and *TCF7L1* was hypermethylated at two CpGs (Additional Fig. 4 E). These CpGs are annotated by ENCODE as a region of an active promoter in embryonic stem cells, indicating a putative regulatory site for the expression of *WNT5A* and *TCF7* in gastric stem cells. In summary, transcriptomic and epigenomic data point towards the implication of the canonical and non-canonical Wnt pathways across stomach regions with differences in strength along an inferior-superior axis.

**Figure 4:**
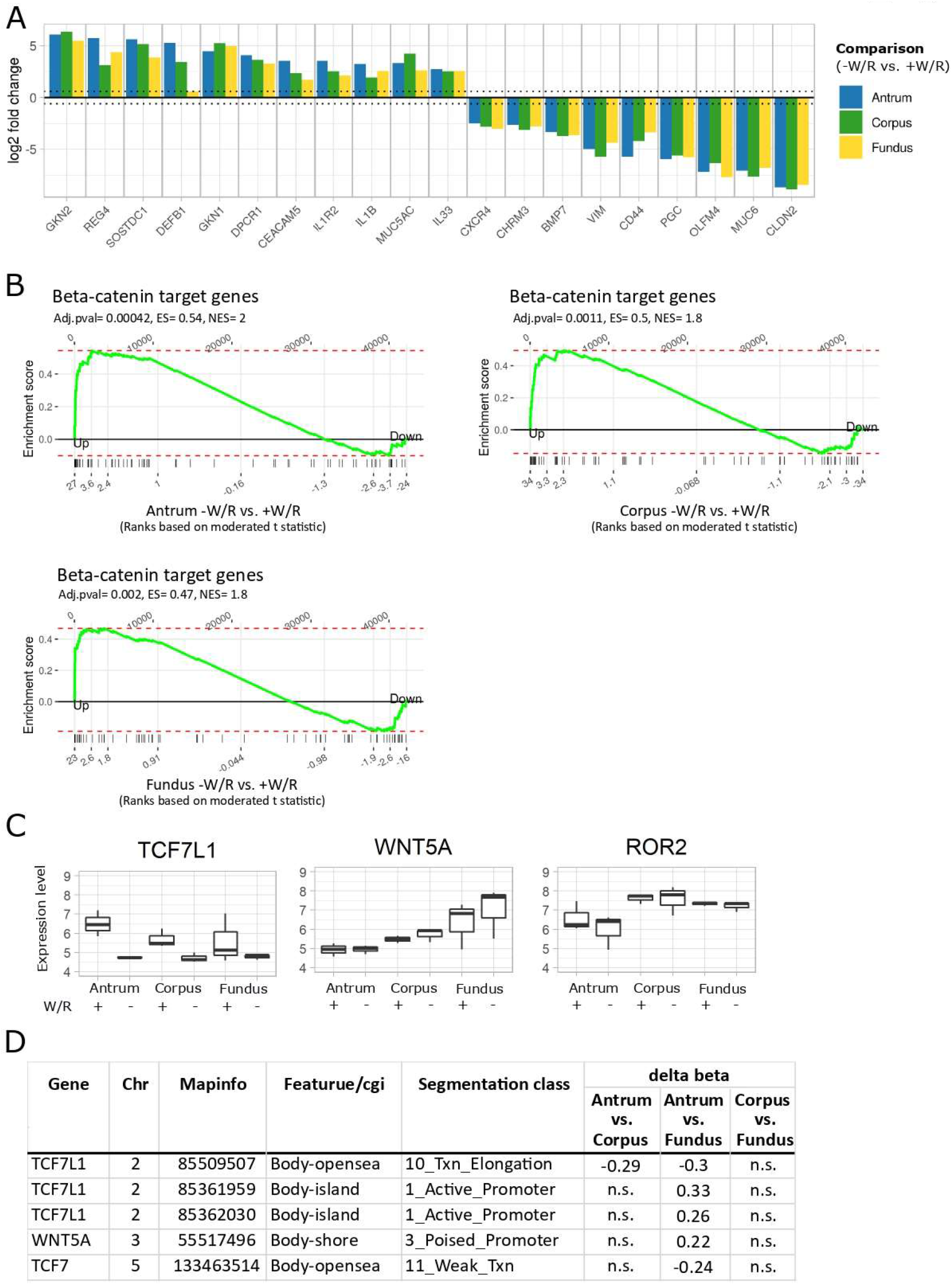
Differentiation- and stem cell-specific gene expression. **A)** Selected up- and down-regulated pit cell- and stem cell-specific genes between -W/R and +W/R conditions in antrum, corpus, and fundus-derived cell cultures (FDR < 0.05) **B)** Gene set enrichment of beta-catenin target genes (n = 66) (Herbst et al., 2014) compared to differential gene expression between -W/R vs. +W/R. The moderated t-score of the DE results was used for ranking the genes. ES - enrichment score; NES - normalized enrichment score. **C)** Normalized gene expression boxplots of *TCF7L1* and the non-canonical Wnt signaling pathway genes *WNT5A* and *ROR2*. The boxplot displays the median, minimum and maximum normalized expression values of three biological replicates. **D)** Table of DM CpGs within the genes displayed in C. n.s. - not significant.

### *Ex vivo*-cultivated organoids of intestinal metaplasia show specific methylation patterns that are maintained in gastric cancer

IM is the emergence of epithelial cells with an at least partial intestinal phenotype in the stomach, associated with chronic *Helicobacter pylori* infection. We cultured epithelial cells obtained from biopsies of antral IM sites and normal control sites *ex vivo*. As expected, *ex vivo* IM organoids showed transcriptional downregulation of stomach-specific genes and upregulation of intestinal genes (Suppl Fig. 5A). These organoids were also clearly segregated from their healthy controls in terms of DNA methylation (Additional Fig. 5B). Additional samples that we cultivated from atrophic mucosa biopsies showed an intermediate methylation phenotype and were not further characterized. The differences detected in IM compared to normal *ex vivo* samples (delta beta > 0.2 or < −0.2, p < 0.05) correlated strongly with differences *in vivo* (Additional Fig. 5C-E), demonstrating the stability of IM epigenomic patterns *ex vivo*. We observed, however, higher DM CpG numbers in the *in vivo* data set compared to the *ex vivo* data set, most likely due to the higher power to detect DM in larger cohorts. We also observed a strong bias of hypermethylation in the *in vivo* data set (14,023 hyper- and 1,199 hypomethylated CpGs, Additional Fig. 5E, left). Even stronger overlaps with our data, including affected genes, promoters and enhancers at the CpG level, were found when selecting the subset of *in vivo* samples with the highest global methylation levels and a higher purity (Huang et al., 2018) (Additional Fig. 5E, right). Higher levels of intermediate DNA methylation due to contaminating cells and proportionally more DM sites have been previously described when comparing results obtained with pure epithelial cells compared to biopsies (Barnicle et al., 2016). Here, we show that the analysis of *ex vivo* cultured gastric cells results in lower admixture effects and improved calling of DM sites (Additional Fig. 5F).

The number of DM CpGs in IM *ex vivo* is similar to what we observed inter-regionally; however, these DM CpGs affected much higher numbers of genes and promoters (Additional Fig. 2A and 5E). Unexpectedly though, only a small number of genes with aberrant promoter hypermethylation or hypomethylation showed differences in gene expression (15/191 and 13/85, respectively; Table 1-2). Nevertheless, these differences were aligned with the general dogma of promoter hypomethylation, leading to increased transcription and *vice versa*. Hypomethylated and upregulated genes included the intestinal stem cell marker Achaete-Scute Family BHLH Transcription Factor 2 (*ASCL2*) as well as two other intestine-specific genes, Tripartite Motif Containing 15 (*TRIM15*) and Fucosyltransferase 6 (*FUT6*) (Uhlén et al., 2015). To further support the regulation of these intestinal genes by promoter methylation, we treated healthy antral mucosoids with the demethylating agent 5-aza2’-deoxycytidine (5aza); this resulted in an increased expression (Table 2, Additional Table 6). We asked if hypomethylated CpGs in *ex vivo* IM also played a possible role in gene deregulation in IM. Enrichment analysis of those DM CpGs revealed enriched binding sites for the intestinal TFs HNF1 Homeobox A (HNF1A), Interferon Regulatory Factor 1 (IRF1), PDX1, and Homeobox A5 (HOXA5) (Fig. 5A). *PDX1* and *HOXA5 gene* expression was increased in *ex vivo*-cultivated organoids of IM (log2 fold change = 1.06 and 0.54, and FDR = 0.174 and 0.025, respectively). CDX2 is the major TF driving intestinal transcriptional program and is known to be upregulated in IM (Barros et al., 2012; Boyd et al., 2010). It has been reported that promoter methylation is not associated with *CDX2* expression in IM tissue (Pereira et al., 2009); the sites of regulation within *CDX2* that provoke its aberrant expression in IM remain unknown. In our IM analysis, we observed accompanying *CDX2* upregulation, promoter hypermethylation as well as a hypomethylation site in the *CDX2* gene body. This gene body site, which contains an EZH2 binding site (Additional Table 4), was also DM in a comparison between the healthy antrum and the corpus/fundus (Fig. 2A), indicating its putative regulatory role (Fig. 5B). Taking into account the *in vivo* data set, we noticed that the DM of this *CDX2* site might be obscured by the admixture effect described above (Additional Fig. 5F). As observed by Huang *et al*. (Huang et al., 2018), we observed an enrichment of DM CpGs in *ex vivo* IM at genes with a bivalent promoter state indicated by the enrichment of H3K4me1 and H3K27me3 (Rada-Iglesias et al., 2011) and TFBS of SUZ12 and EZH2 (Additional Fig. 6A). Additionally, in our data also hypomethylated CpGs show this enrichment, probably because we detected higher numbers of hypomethylated CpGs. Together, these data indicate that both key genes and TF binding sites might be affected by DM patterns in IM.

**Figure 5:**
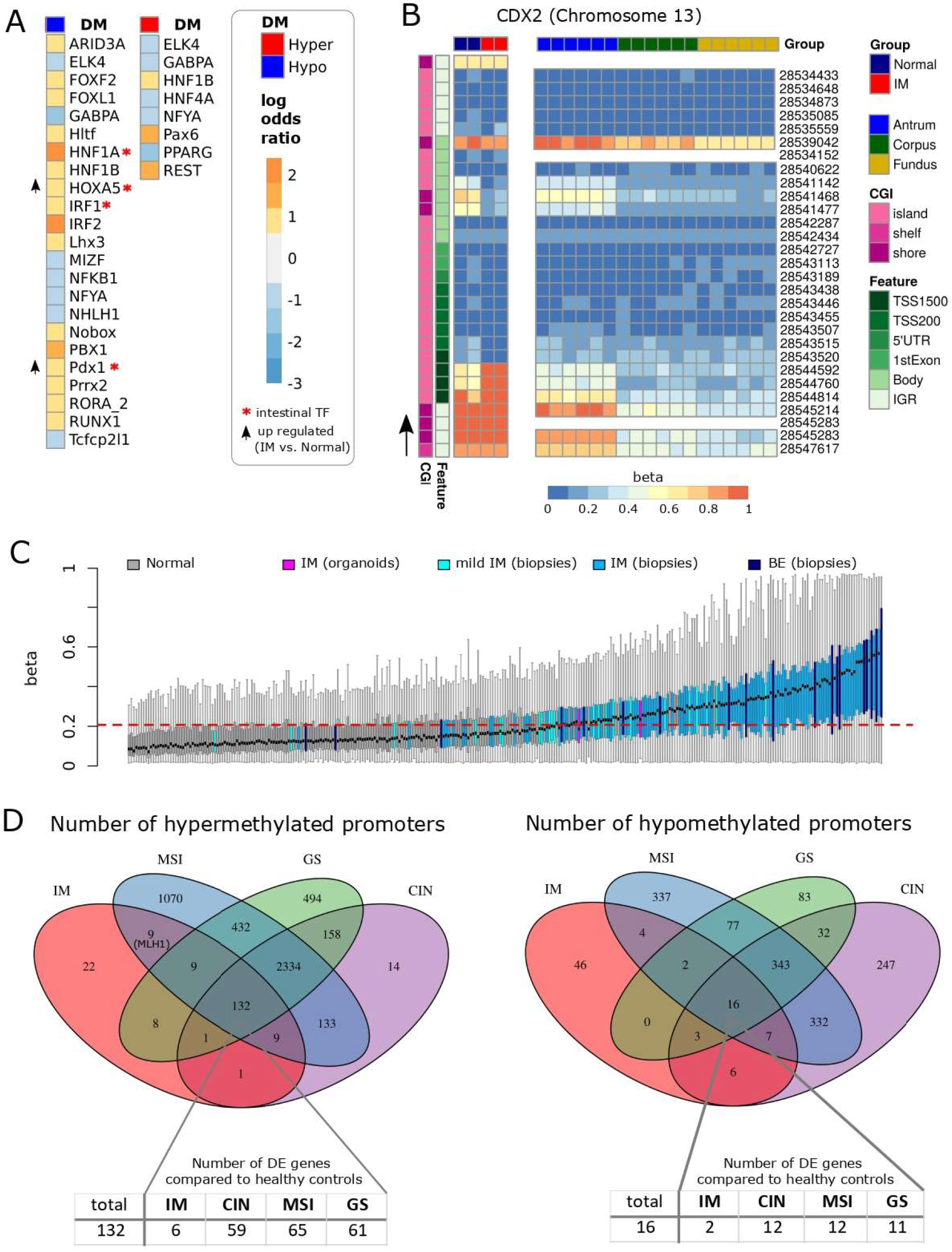
Differential methylation in intestinal metaplasia. **A)** Enrichment of hypermethylated and hypomethylated CpGs in TF binding sites (JASPAR collection) in organoids derived from intestinal metaplasia (IM) biopsies compared to organoids derived from normal gastric biopsies. Shown are log odds ratios of significantly enriched (log odds ratio > 0.6) or depleted (log odds ratio < −0.6) genomic features (FDR < 0.05). Color code ranges from blue (depleted) to red (enriched); grey refers to a log odds ratio between −0.6 and −0.6. Hyper - hypermethylated CpGs; Hypo - hypomethylated CpGs; CpG island (CGI). **B)** Heatmap of the *CDX2* DNA methylation pattern in the IM organoids vs. normal gastric organoids. The arrow indicates the transcriptional direction. **C)** ExE hyper CGI methylation of in healthy and precancerous samples. Shown are the mean ExE-hyper-CpG island methylation values per sample ranked by their means. Normal samples include all the samples from our mucosoid cultures (antrum, corpus, and fundus) as well as all the healthy samples from the organoid and biopsy IM data sets (Huang et al., 2018) and the Barrett’s esophagus (BE) biopsy data sets (Krause et al., 2016). The dashed red line indicates the upper 95 % confidence interval limit of all normal samples. Diseased samples include the respective *in vivo* mild IM and IM (biopsies), IM (organoids), and BE (biopsies). **D)** Venn diagram of genes with hypermethylated or hypomethylated promoter regions in diseased samples as compared to healthy samples. Diseased samples: IM (organoids), microsatellite instable (MSI), genomically stable (GS), and chromosomal instable (CIN) molecular subtypes of gastric cancer (Cancer Genome Atlas Research Network, 2014). Tables indicate the number of differentially expressed genes among those shared by all three cancer subtypes and IM.

**Table 1:** Hypermethylated promoter regions in IM.

| Gene | Hypermethylated CpGs/Promoter region | Total CpGs/promoter | logFC | p-value | FDR | detected in |
| --- | --- | --- | --- | --- | --- | --- |
| KDR | 3 | 12 | -1.5 | 0.0000 | 0.0003 | Ex vivo and in vivo |
| GCNT2 | 3 | 17 | -1.42 | 0.0000 | 0.0000 | Ex vivo |
| CAPN13 | 3 | 6 | -1.39 | 0.0000 | 0.0000 | Ex vivo |
| EYA4 | 7 | 51 | -1.37 | 0.0000 | 0.0013 | Ex vivo and in vivo |
| ITGBL1 | 4 | 10 | -1.22 | 0.0001 | 0.0101 | Ex vivo |
| SLC9A3 | 5 | 7 | -1.01 | 0.0001 | 0.0112 | Ex vivo |
| CLIC6 | 5 | 11 | -0.89 | 0.0039 | 0.1244 | Ex vivo |
| CHAD | 7 | 8 | -0.69 | 0.0312 | 0.4136 | Ex vivo |
| SORBS2 | 7 | 55 | -0.69 | 0.0003 | 0.0201 | Ex vivo |
| TAS1R1 | 3 | 14 | -0.68 | 0.0293 | 0.4018 | Ex vivo |
| PSCA | 4 | 6 | -0.66 | 0.0266 | 0.3812 | Ex vivo |
| CYP2E1 | 5 | 7 | 0.77 | 0.0063 | 0.1681 | Ex vivo |
| TMEM176B | 2 | 20 | 0.89 | 0.0022 | 0.0838 | Ex vivo |
| GJB2 | 3 | 20 | 1.05 | 0.0000 | 0.0021 | Ex vivo |
| CDX2 | 2 | 12 | 1.34 | 0.0000 | 0.0033 | Ex vivo |

Shown are the 15 genes with hypermethylated promoter regions in IM that also show altered gene expression. The *in vivo* samples represent antral samples with a high methylation cluster. logFC = log 2 fold change; FDR = false discovery rate.

**Table 2:** Hypomethylated promoter regions in IM.

| gene | Hypomethylated<br>CpGs/Promoter<br>region | Total CpGs/<br>promoter | logFC | p-value | FDR | detected in | 5aza<br>logFC |
| --- | --- | --- | --- | --- | --- | --- | --- |
| **TRIM15** | 9 | 25 | 2.11 | 0.0000 | 0.0000 | Ex vivo and in<br>vivo | 1.66 |
| **ASCL2** | 6 | 45 | 1.27 | 0.0000 | 0.0003 | Ex vivo and in<br>vivo | 0.79 |
| **HOXA2** | 15 | 28 | 1.24 | 0.0000 | 0.0053 | Ex vivo |  |
| **HMGA1** | 3 | 17 | 1.09 | 0.0000 | 0.0000 | Ex vivo |  |
| **TMEM154** | 3 | 7 | 0.92 | 0.0040 | 0.1257 | Ex vivo |  |
| **KRT23** | 2 | 6 | 0.91 | 0.0045 | 0.1351 | Ex vivo | 4.25 |
| **FUT6** | 4 | 8 | 0.90 | 0.0033 | 0.1117 | Ex vivo and in<br>vivo | 0.99 |
| **GAL3ST2** | 2 | 6 | 0.88 | 0.0006 | 0.0315 | Ex vivo and in<br>vivo |  |
| **GJC2** | 2 | 13 | 0.67 | 0.0370 | 0.4529 | Ex vivo | 2.89 |
| **SYNE1** | 2 | 21 | 0.66 | 0.0266 | 0.3812 | Ex vivo | 1.48 |
| **C8orf74** | 2 | 6 | 0.64 | 0.0469 | 0.5108 | Ex vivo |  |
| **HOXA4** | 10 | 24 | 0.62 | 0.0262 | 0.3786 | Ex vivo |  |
| **DCHS2** | 2 | 15 | -0.64 | 0.0093 | 0.2145 | Ex vivo |  |

Shown are the 13 genes with hypomethylated promoter regions in IM that also show altered gene expression. The in vivo samples represent antral samples with a high methylation cluster. 5 aza = 5- aza-2'-deoxycytidine; logFC = log 2 fold change; FDR = false discovery rate.

We were interested in understanding the mechanisms that drive aberrant methylation in IM and its relevance to GC. Recently, a hypermethylation signature at specific CpG islands shared by most cancers and associated with extra-embryonic tissues was identified (Smith et al., 2017). Interestingly, IM organoids displayed increased methylation levels at these sites compared to normal tissues. (Fig. 5C, Additional Fig. 6B-C). We observed the same pattern in the *in vivo* data set of IM (Huang et al., 2018) as well as in a data set of Barrett’s esophagus (BE) samples (Krause et al., 2016), which is an IM that occurs in the lower esophagus. These results suggest that the dysregulation of putative responsible signaling pathways occurs already before the transformation. Finally, we asked if any IM-related changes are preserved in GC. We determined the overlaps of hypermethylated and hypomethylated promoters in IM with those identified in molecular subtypes of GC, namely chromosomally instable (CIN), microsatellite instable (MSI) and genomically stable (GS) GCs. Most intestinal-like GCs (59 %) are classified as belonging to the CIN subtype, whereas MSI and GS subtypes represent only 24 and 9 % of intestinal-like GCs, respectively (Cancer Genome Atlas Research Network, 2014). A recent study (Huang et al., 2018) has found a substantial overlap between hypermethylated regions in IM and those of CIN and MSI subtypes of GC. Our data is in good agreement with these findings. We found that 69 % of the hypermethylated promoters and 19% of the hypomethylated promoters in IM overlapped with all three molecular GC subtypes (Fig. 5D). Since aberrant promoter methylation in IM was shared with all GC subtypes, it seems that these differences might not be limited to the intestinal phenotype alone. We further observed that while most genes with shared aberrant promoter methylation in IM and GC are not differentially expressed in IM, they show deregulation in cancer samples compared to healthy controls (Fig. 5D, lower row). This indicates that nascent aberrant methylation of these genes does not lead to considerable regulatory effects at the IM stage, but it does once the cells become malignant. As a prominent example, promoter hypermethylation of MutL Homolog 1 (*MLH1*) is present already in IM, but the hypermethylation does not affect the specific CpG island responsible for its silencing in cancer (Morak et al., 2008) (Additional Fig. 6D); *MLH1* was not differentially expressed (data not shown). These data support the hypothesis that aberrant methylation starting in IM continues in GC, eventually promoting deregulated expression.

## Discussion

The stomach, a major organ of the gastrointestinal tract, is divided into several functionally distinct domains along its longitudinal (craniocaudal) axis; these domains are characterized by phenotypically and functionally diverse mucosa with the predominance of distinct cell types and lineages. While some regulatory factors of the different zones have been identified (McCracken et al., 2014), it is unclear what their role is in inter-regional stomach development and how these phenotypic differences are maintained. DNA methylation as a stable and inheritable epigenetic modification is likely to play a role in region-specific cell differentiation.

By using a primary human gastric epithelial cell culture model (Boccellato et al., 2018), we show that global DNA methylation patterns define a regional identity that clearly and stably distinguishes the antrum, corpus, and fundus. We present an extensive analysis across all three main stomach regions and differentiation states (stem and pit cells), thereby creating a whole map of DNA methylation and gene expression dynamics in the healthy human gastric epithelium. We present novel evidence of a longitudinal patterning of epigenetic and transcriptional markers in the stomach, involving developmentally important TFs and major pathways that control stemness and differentiation. Interestingly, we find that many DM TFs are no longer transcriptionally active in the adult stomach and that most region-specific lineage genes are not affected by DM. Together, these results infer plasticity of gastric epithelial stem cells and a specification of regional cellular phenotypes by only a few TFs and possibly cell-extrinsic signals. Our results are supported by previous studies that have shown region-specific methylation profiles in individual tissues (Irizarry et al., 2009; Ziller et al., 2013) and different colon regions (Barnicle et al., 2016). Similarly, we observed inter-regional DM of homeobox genes, while stomach-specific genes were rarely affected by DM. Moreover, we found that significant changes in DNA methylation do not accompany the differentiation of stem cell-enriched gastric mucosoids to pit cells; this has been similarly observed in primary intestinal epithelial cells (Kraiczy et al., 2019). In addition to TFs, such as PDX1 and IRX5, known to be involved in stomach development and region specification, we identified several novel TFs, including SIM2, MEIS1, MEIS2, and CDX2. Mechanistically, most of those genes were DM at enhancers and promoters that bind EZH2 and SUZ12, pointing towards their possible regulation by PCR2. For many of those factors, we could validate DM and partial differential expression by comparing stomach regions across several stomach datasets, thus supporting their potential role in the induction of region-specific differentiation. MEIS TFs can interact with other homeodomain TFs alone or in complexes to fine-tune their DNA binding characteristics (Schulte and Geerts, 2019). Interestingly, we identified DM in one of their binding partners, PDX1, which is required for antral G-cell differentiation (Larsson et al., 1996). In different published *in vivo* data sets as well as in our *in vitro* data set, MEIS1 and MEIS2 showed higher expression in the antrum than the corpus, whereas SIM2 showed higher expression in the corpus than the antrum. The role of SIM2 in the stomach is yet to be investigated. Notably, in the small intestine, SIM2 has been shown to activate the transcription of the Wnt signaling mediator *TCF7L2* and to directly regulate the expression of several antimicrobial peptides in the small intestine (Chen et al., 2014). Furthermore, we provide evidence for the low expression of CDX2 in the antrum compared to its strict suppression in the proximal stomach. In the human stomach, the corpus and fundus are often referred to synonymously as they functionally share the characteristic chief- and parietal cells (McCracken et al., 2017). However, even in the absence of these cell types in our analyzed cell culture model, we observed unique features of the most proximal stomach region that are absent in the corpus, such as high levels of complement and immune response genes and the lack of *LGR5* expression.

Wnt and R-Spondin are major stemness regulators in the intestine and stomach (Barker et al., 2007; Fischer and Sigal, 2019). *Lgr5,* a main component of the Wnt pathway, is usually expressed in the antrum, while the mouse corpus usually lacks *Lgr5* (Barker et al., 2010), suggesting different roles of the Wnt pathway in different stomach regions. In contrast, +W/R conditions led to comparable activation of beta-catenin signaling and maintained a stem-like phenotype in epithelial cells *ex-vivo* independently of origin. Yet, the expression of the Wnt-pathway mediator *TCF7L1* was attenuated in the proximal stomach, while major effectors of the antagonistic non-canonical Wnt Signaling, *ROR2* and *WNT5A* (Miyoshi et al., 2012), were enhanced compared to the inferior regions. These findings, which indicate differential Wnt signaling in the superior stomach, are supported by DM in the promoter and enhancer sites of *TCF7L1*. Such region-specific regulation of differentiation-associated genes has been observed in intestinal organoids as well (Kraiczy et al., 2019). In agreement with our findings, a supportive role for non-canonical *Wnt5a* signaling in the stem cell niche of the gastric corpus in contrast to that of the antrum has been suggested by Hayakawa et al. (2015). The expression of *WNT5A* and *ROR2* in the human corpus and fundus, as detected by us, indicates that gastric epithelial cells of the fundic gland are susceptible to WNT5A signaling. It remains to be determined whether Wnt signaling contributes to the stem cell niche as proposed by Hayakawa et al. (2015).

We extended our analysis of epithelial cell-specific DNA methylation in an attempt to characterize the differences in diseased primary epithelial cells. Comparing healthy samples to GC and its precancerous lesion IM suggested an early occurrence of GC-specific hypermethylation and hypomethylation patterns already at the IM stage. First, we observed hypermethylation in *ex vivo*-cultivated IM as well as in samples of IM *in vivo* and BE. This hypermethylation signature affects a set of particular CpG islands, related to extra-embryonic ectoderm-specific genes found in most cancer types, and thought to result from dysregulated Fibroblast Growth Factor 2 (FGF2) signaling (Smith et al., 2017). Although we did not detect differential expression of FGF pathway genes in IM *ex vivo* (data not shown), the injection of the cytotoxin-associated gene A (CagA) by *Helicobacter pylori* is known to activate FGF signaling (Katoh and Katoh, 2006). Similarly, dysregulated FGF signaling has been observed in BE (Barclay et al., 2005; Kresty et al., 2011). Second, similar to Huang and others, we observed this shared hypermethylation pattern by comparing IM *ex vivo* and GC subtypes. Most likely, due to our model of pure epithelial cells, we were also able to identify several hypomethylated promoters that are shared between IM and all GC subtypes, excluding the EBV subtype (Cancer Genome Atlas Research Network, 2014). We further focused on the shared genes with aberrant hypermethylated or hypomethylated promoters. We realized that their corresponding gene expression differences, compared to healthy controls, almost exclusively occurred in GC samples but not in IM. This indicates that additional (epi-) genetic changes must occur during the progression of the pre-neoplastic lesions towards full-blown GC.

While the *ex vivo*-cultivated IM organoids correlated well with those differences *in vivo*, the lack of cell admixture in cultured cells (Barnicle et al., 2016) allowed more sensitive detection of hypomethylated CpGs sites in IM as compared to studies involving biopsies (Huang et al., 2018). Thus, we found hypomethylation of the promoter region and upregulation of the putative stem cell marker of the small intestine, *ASCL2*, essential for maintaining adult intestinal stem cells (van der Flier et al., 2009). In addition to the induction of intestinal genes, we also found that hypomethylated CpGs were enriched at binding sites of intestinal TFs, such as HNF1A, HOXA5, IRF2 and PDX1. These intestinal TFs were partially upregulated in IM *ex vivo*, compatible with the altered epigenetic regulation of transcriptional networks in IM. Both the binding of TFs at hypomethylated sites (Domcke et al., 2015) and TF binding-induced hypomethylation (Stadler et al., 2011) have been previously observed. Further, enhancer hypomethylation in the healthy intestine has been shown to be associated with inappropriate TF binding (Sheaffer et al., 2014). Therefore, our results reveal several potential epigenetic mechanisms and target genes that could contribute to molecular changes that promote carcinogenesis in gastric epithelial cells.

### Conclusions

In summary, we present an extensive characterization of epigenetic and transcriptional landscapes that vary across stomach regions. Our study reveals new candidate regulators for region-specific phenotypes and illuminates crucial mechanisms of cellular transformation.

## Methods

### Human Material

#### Gastric sleeve

Human gastric tissue samples were obtained from the Center for Bariatric and Metabolic Surgery at the Charité University Medicine, Berlin, Germany. Patient samples, negative for *Helicobacter pylori*, were used for isolation of gastric glands from the antrum, corpus, and fundus.

#### Gastric biopsy

Human gastric biopsies of antral normal, atrophic, and metaplastic gastric tissue were provided by the Department of Gastroenterology, Hepatology and Infectious Diseases, Otto-von-Guericke-University of Magdeburg, Magdeburg, Germany. Adjacent biopsies were assessed macro-and microscopically by a pathologist, and samples, where macro-and microscopic assessment matched, were used for the isolation of epithelial cells.

### Primary cell culture

Organoids and Mucosoids were cultivated as previously described (Boccellato et al., 2018; Schlaermann et al., 2016). Primary epithelial cells from the corpus and fundus were cultivated under the same conditions as the antral mucosoids. Differentiation of the mucosoids was achieved by replacing Wnt and R-spondin in the cell culture medium with advanced Dulbecco’s modified Eagle medium/F12 (Invitrogen) for seven days. Samples of undifferentiated antrum and corpus have also been used for gene expression analysis by Wölffling *et al*. (Wölffling et al., 2021).

### 5-aza-2’-deoxycytidine treatment of mucosoid cultures

Freshly-seeded healthy antral undifferentiated mucosoid cultures were treated every 24 h with 4 µM 5aza (in 0.02 % acetic acid; Sigma-Aldrich). A global decrease in DNA methylation, measured with the 5mC ELISA kit (Zymo Research), to 12 % of the total human 5-methylated cytosine (5mC) content was achieved after 10 days, after which cells were harvested for RNA isolation.

### Nucleic Acid isolation

DNA and RNA were extracted from mucosoid cultures using the QIAamp DNA Mini Kit (Qiagen) and Trizol (Thermo Fisher Scientific) methods, respectively. To extract DNA and RNA from *ex vivo*-cultivated organoids and 5aza-treated mucosoids, we used the AllPrep DNA/RNA/miRNA Universal Kit (Qiagen). RNA integrity and quantity of RNA were assessed using the Agilent 2100 Bioanalyzer (Agilent). DNA concentration was measured using the Qubit dsDNA HS Assay Kit (Thermo Fisher Scientific), and integrity was assessed by gel electrophoresis (0.8 % agarose gel, 120 V, 1 h).

### DNA methylation array (450k and EPIC)

The Illumina Infinium® human 450k (Illumina, WG-314-10031) and EPIC methylation (Illumina, SWG-317-1001) bead chips were applied for *ex vivo*-cultivated organoids and healthy mucosoids, respectively, according to the manufacturer’s protocol at the Life&Brain research center (Bonn, Germany) and in collaboration with Per Hoffmann from the Institute of Human Genetics, University of Bonn.

### Gene expression microarray

Single color Agilent SurePrint G3 Custom Gene Expression Microarray 8 x 60K (Agilent, Agilent-048908; GEO Platform GPL21272) was used for gene expression analysis according to the manufacturer’s protocol. Note: a part of the gene expression results obtained with the samples of +W/R antrum and +/W/R corpus has already been published (Wölffling et al., 2021). Here, we used these samples combined with the respective -W/R samples and extended the data set to +W/R and -W/R fundus.

### Bioinformatic methods

Data analysis was performed in the R/Bioconductor environment (R Core Team (2017), n.d.). Public data sets used in this work were downloaded from GEO (Additional table 7), TCGA (STAD) (Cancer Genome Atlas Research Network, 2014), GTEx Portal (GTEx Consortium et al., 2017) and the Human Protein Atlas (Uhlén et al., 2015).

### DNA methylation

Raw intensity files (idats) were processed using the ChAMP package (Morris et al., 2014; Tian et al., 2017) using default settings. When data sets of the same platform were combined, all idat files were loaded and processed together. Only when combining 450k and EPIC data sets, the minfi (Tab. 2.16) function combineArrays() was used to virtually combine the already loaded and filtered 450k and EPIC objects (Aryee et al., 2014; Fortin et al., 2017). Subset-quantile Within Array Normalization (SWAN) was used to create data sets in this study. For combined data sets, functional normalization (Fortin et al., 2014) was applied. Patient and slide effects were adjusted in the healthy *in vitro* stomach data set using the champ.SVD function. Differential methylation analysis was performed with limma (Ritchie et al., 2015), applying a moderated t-test for the mucosoids and a paired moderated t-test for the *ex vivo* data set using M-values. Since beta-values, ranging from 0 (unmethylated) to 1 (methylated), are more intuitive, DM in this manuscript is reported as delta beta. For the regional comparisons, +W/R and - W/R samples of the same region were combined and analyzed as six biological replicates. Genes were considered DM when at least two CpGs/gene were affected by significant differential methylation (FDR < 0.05, delta beta > 20 %). The threshold was lowered to 1 CpG/gene when Wnt pathway genes were determined as DM. Promoters were considered DM when at least one CpG/promoter region, defined as TSS1500, TSS200, 5’UTR or 1st Exon, was significantly differentially methylated. Reference for genes of the canonical and non-canonical Wnt pathways was taken from the GO-Wnt and GO-non-canonical-Wnt gene lists. Stomach-specific genes were taken from the Human Protein Atlas as those that show five-fold higher mRNA levels compared to average levels in all other tissues (Uhlén et al., 2015). Chief and parietal cell-specific genes were determined from re-analyzing the data set of Ramsey and others (Ramsey et al., 2007), who profiled murine gastric epithelial cells, as compared to the other cell types. Symbols of mouse genes whose expression was logFC > 1 were translated into human gene symbols and used as a gene set. Genes of the Troy+ cell signature were taken from the Additional Table 1 (Stange et al., 2013). DM genes were classified as TFs according to (Vaquerizas et al., 2009). Protein-protein interactions and enrichment in GO-biological processes of DM genes were determined using the platform StringDB (Szklarczyk et al., 2015). The segmentation classes for specific gene annotations were taken from the UCSC table browser bed files of Broad Hmm in H1hesc for table annotation and the average segmentation combined H1hesc for visualization. CpG sets corresponding to hypermethylated or hypomethylated CpGs were created from DM results. Only CpGs with delta beta > 20 % were included to reduce the maximum size of the CpG set to < 3,000; 5,000 permutations were applied. The distribution of DM CpGs along genomic features was determined using the UCSC RefGene Group from the Illumina annotation of the EPIC or 450k array by calculating Pearson residuals of DM CpGs for each genomic feature and CpG island relation. Enrichment of DM CpGs was determined using Locus overlap analysis (LOLA) (Sheffield and Bock, 2016), the ENCODE Segmentation classes and TFBSs (Siggens and Ekwall, 2014), the UCSC features (Kent et al., 2002), the Roadmap epigenomics histone marks (Roadmap Epigenomics Consortium et al., 2015) and the JASPAR motifs collection (Khan et al., 2018). For all core databases, only the data of untreated embryonic cell lines were included. The Roadmap epigenomics collection was subsetted for tissues corresponding to the respective comparisons. For the inter-regional comparison in the stomach, histone marks determined in the fetal stomach, stomach mucosa, gastric and stomach smooth muscle are included. For the comparison between tissues, we additionally included data from the fetal small and large intestine, small intestine, sigmoid colon, colonic mucosa, duodenum mucosa and esophagus. The function fisher.test() within the runLOLA() function was modified in order to allow a two-sided Fisher’s exact test. We determined the high methylation cluster in samples of IM *in vivo* (Huang et al., 2018) by performing hierarchical clustering of the 108 IM samples and selecting the 39 antral samples with the high methylation cluster. Extra-embryonic ectoderm (ExE)-hypermethylated CpG islands (Smith et al., 2017) were orthologously mapped to the human hg19 genome using the UCSC Genome Browser liftOver tool (Kent et al., 2002). Bed files of the genomic regions were mapped to the closest human CpG island, and the mean methylation/CpG island was determined. The mucosoid, the *in vivo* IM samples, and the BE data set included 483 of the 489 mouse CpG islands, whereas in the *ex vivo* IM data set 487 CpG islands were represented. Significance between diseased and normal samples was determined by a Wilcoxon rank-sum test with continuity correction between the mean methylation of all ExE hyper CpG islands in diseased data sets compared to all normal means. Healthy samples from the Roadmap genome collection that were included in the Smith publication were included as well. The distribution of average methylation of all other CpG island promoters, which were interrogated on the methylation arrays, and of the ExE hyper CpG islands was assigned as a control. Hypermethylated sites of *MLH1* in cancer were taken from (Morak et al., 2008), using also the UCSC genome browser function lift over tool to visualize the hg18 annotated sites in hg19 with custom tracks of DM CpGs *ex vivo* (Kent et al., 2002).

### Gene expression

Background correction was performed using the normexp-method (Ritchie et al., 2007), and inter-array normalization was performed with the quantile method of (Bolstad et al., 2003). Differential gene expression was determined using limma by applying a moderated paired t-test. Gene set enrichments included gene sets with sizes between 15 and 2,000 genes; 5,000 permutations were applied. The beta-catenin gene set was taken from (Herbst et al., 2014). Gene set enrichment analysis was performed using the fgsea package and gene sets of hallmark, pathway, motif, GO_BP, Oncogenic, and immunologic of the Molecular Signature DataBase (Subramanian et al., 2005). RNA-Seq-based gene expression values (RSEM FPKM) from normal stomach samples of the GTEX project and stomach adenocarcinoma samples of the TCGA project were obtained from the TOIL project website (https://xenabrowser.net/datapages/?cohort=TCGA%20TARGET%20GTEx) and used to identify gene expression changes in DM genes in GC (Vivian et al., 2017).

## Declarations

### Ethics approval

The Charité University Medicine ethics committee approved the scientific usage of gastric sleeve resection material (EA1/129/12), and pseudonymized samples from patients. The Otto-von-Guericke-University of Magdeburg ethics committee approved the scientific usage of human gastric biopsies (EA 80/11).

### Data Availability

The DNA methylation and gene expression datasets generated during the current study have been deposited in the National Centre for Biotechnology Information Omnibus (GEO) under accession code GSE141660. Reviewer access token: mtivkcsallcdfsb.

### Competing Interests

The authors declare that they have no competing interests.

### Funding

This work was supported by the European Research Council (ERC) Advanced Grant (885008-MADMICS) and the European Sequencing and Genotyping Infrastructure (ESGI) grant (262055-BIGC) to TFM.

### Author contributions

KF, FB, HB and TFM conceived the project and designed experiments, KF performed air liquid culture experiments of stomach samples, KF. performed methylation analysis, PS and MK performed organoid cultures of ex vivo stomach samples, CD selected and provided gastric samples, KF and HB performed the bioinformatic analysis. IG generated and provided RNA-Seq data for ex-vivo samples. KF, HB and TFM wrote the manuscript.

## Acknowledgements

We thank Kfir Lapid for editing the manuscript. We are also grateful to P. Hoffmann, T. Kätzel, and S. Herms, Institute of Human Genetics, University of Bonn, for performing the methylation arrays and initial support in raw data processing. We further thank I. Wagner and H. Mollenkopf, Max Planck Institute for Infection Biology, Berlin, Germany, for performing the microarrays.

## Notes

https://www.ncbi.nlm.nih.gov/geo/query/acc.cgi?acc=GSE141660

### Competing Interest Statement

The authors have declared no competing interest.

## References

Arnold, K., Sarkar, A., Yram, M.A., Polo, J.M., Bronson, R., Sengupta, S., Seandel, M., Geijsen, N., Hochedlinger, K., 2011. Sox2(+) adult stem and progenitor cells are important for tissue regeneration and survival of mice. Cell Stem Cell 9, 317–329. https://doi.org/10.1016/j.stem.2011.09.001

Aryee, M.J., Jaffe, A.E., Corrada-Bravo, H., Ladd-Acosta, C., Feinberg, A.P., Hansen, K.D., Irizarry, R.A., 2014. Minfi: a flexible and comprehensive Bioconductor package for the analysis of Infinium DNA methylation microarrays. Bioinformatics 30, 1363–1369. https://doi.org/10.1093/bioinformatics/btu049

Barker, N., Huch, M., Kujala, P., van de Wetering, M., Snippert, H.J., van Es, J.H., Sato, T., Stange, D.E., Begthel, H., van den Born, M., Danenberg, E., van den Brink, S., Korving, J., Abo, A., Peters, P.J., Wright, N., Poulsom, R., Clevers, H., 2010. Lgr5(+ve) stem cells drive self-renewal in the stomach and build long-lived gastric units in vitro. Cell Stem Cell 6, 25–36. https://doi.org/10.1016/j.stem.2009.11.013

Barker, N., van Es, J.H., Kuipers, J., Kujala, P., van den Born, M., Cozijnsen, M., Haegebarth, A., Korving, J., Begthel, H., Peters, P.J., Clevers, H., 2007. Identification of stem cells in small intestine and colon by marker gene Lgr5. Nature 449, 1003–1007. https://doi.org/10.1038/nature06196

Barnicle, A., Seoighe, C., Golden, A., Greally, J.M., Egan, L.J., 2016. Differential DNA methylation patterns of homeobox genes in proximal and distal colon epithelial cells. Physiol. Genomics 48, 257–273. https://doi.org/10.1152/physiolgenomics.00046.2015

Barros, R., Freund, J.-N., David, L., Almeida, R., 2012. Gastric intestinal metaplasia revisited: function and regulation of CDX2. Trends in Molecular Medicine 18, 555–563. https://doi.org/10.1016/j.molmed.2012.07.006

Bartfeld, S., Bayram, T., van de Wetering, M., Huch, M., Begthel, H., Kujala, P., Vries, R., Peters, P.J., Clevers, H., 2015. In vitro expansion of human gastric epithelial stem cells and their responses to bacterial infection. Gastroenterology 148, 126–136.e6. https://doi.org/10.1053/j.gastro.2014.09.042

Batool, A., Jin, C., Liu, Y.-X., 2019. Role of EZH2 in cell lineage determination and relative signaling pathways. Front Biosci (Landmark Ed) 24, 947–960.

Beerman, I., Bock, C., Garrison, B.S., Smith, Z.D., Gu, H., Meissner, A., Rossi, D.J., 2013. Proliferation-dependent alterations of the DNA methylation landscape underlie hematopoietic stem cell aging. Cell Stem Cell 12, 413–425. https://doi.org/10.1016/j.stem.2013.01.017

Bibikova, M., Barnes, B., Tsan, C., Ho, V., Klotzle, B., Le, J.M., Delano, D., Zhang, L., Schroth, G.P., Gunderson, K.L., Fan, J.-B., Shen, R., 2011. High density DNA methylation array with single CpG site resolution. Genomics 98, 288–295. https://doi.org/10.1016/j.ygeno.2011.07.007

Bird, A., 2002. DNA methylation patterns and epigenetic memory. Genes Dev. 16, 6–21. https://doi.org/10.1101/gad.947102

Boccellato, F., Woelffling, S., Imai-Matsushima, A., Sanchez, G., Goosmann, C., Schmid, M., Berger, H., Morey, P., Denecke, C., Ordemann, J., Meyer, T.F., 2018. Polarised epithelial monolayers of the gastric mucosa reveal insights into mucosal homeostasis and defence against infection. Gut. https://doi.org/10.1136/gutjnl-2017-314540

Bock, C., Beerman, I., Lien, W.-H., Smith, Z.D., Gu, H., Boyle, P., Gnirke, A., Fuchs, E., Rossi, D.J., Meissner, A., 2012. DNA Methylation Dynamics during In Vivo Differentiation of Blood and Skin Stem Cells. Molecular Cell 47, 633–647. https://doi.org/10.1016/j.molcel.2012.06.019

Bolstad, B.M., Irizarry, R.A., Astrand, M., Speed, T.P., 2003. A comparison of normalization methods for high density oligonucleotide array data based on variance and bias. Bioinformatics 19, 185–193. https://doi.org/10.1093/bioinformatics/19.2.185

Boyd, M., Hansen, M., Jensen, T.G.K., Perearnau, A., Olsen, A.K., Bram, L.L., Bak, M., Tommerup, N., Olsen, J., Troelsen, J.T., 2010. Genome-wide analysis of CDX2 binding in intestinal epithelial cells (Caco-2). J. Biol. Chem. 285, 25115–25125. https://doi.org/10.1074/jbc.M109.089516

Cancer Genome Atlas Research Network, 2014. Comprehensive molecular characterization of gastric adenocarcinoma. Nature 513, 202–209. https://doi.org/10.1038/nature13480

Chen, K.-J., Lizaso, A., Lee, Y.-H., 2014. SIM2 maintains innate host defense of the small intestine. Am J Physiol Gastrointest Liver Physiol 307, G1044–1056. https://doi.org/10.1152/ajpgi.00241.2014

Companioni, O., Sanz-Anquela, J.M., Pardo, M.L., Puigdecanet, E., Nonell, L., García, N., Parra Blanco, V., López, C., Andreu, V., Cuatrecasas, M., Garmendia, M., Gisbert, J.P., Gonzalez, C.A., Sala, N., 2017. Gene expression study and pathway analysis of histological subtypes of intestinal metaplasia that progress to gastric cancer. PLoS One 12, e0176043. https://doi.org/10.1371/journal.pone.0176043

Fischer, A.-S., Sigal, M., 2019. The Role of Wnt and R-spondin in the Stomach During Health and Disease. Biomedicines 7. https://doi.org/10.3390/biomedicines7020044

Fortin, J.-P., Hansen, K.D., 2015. Reconstructing A/B compartments as revealed by Hi-C using long-range correlations in epigenetic data. Genome Biol. 16, 180. https://doi.org/10.1186/s13059-015-0741-y

Fortin, J.-P., Labbe, A., Lemire, M., Zanke, B.W., Hudson, T.J., Fertig, E.J., Greenwood, C.M., Hansen, K.D., 2014. Functional normalization of 450k methylation array data improves replication in large cancer studies. Genome Biol. 15, 503. https://doi.org/10.1186/s13059-014-0503-2

Fortin, J.-P., Triche, T.J., Hansen, K.D., 2017. Preprocessing, normalization and integration of the Illumina HumanMethylationEPIC array with minfi. Bioinformatics 33, 558–560. https://doi.org/10.1093/bioinformatics/btw691

Fouse, S.D., Shen, Y., Pellegrini, M., Cole, S., Meissner, A., Van Neste, L., Jaenisch, R., Fan, G., 2008. Promoter CpG Methylation Contributes to ES Cell Gene Regulation in Parallel with Oct4/Nanog, PcG Complex, and Histone H3 K4/K27 Trimethylation. Cell Stem Cell 2, 160–169. https://doi.org/10.1016/j.stem.2007.12.011

Greenberg, M.V.C., Bourc’his, D., 2019. The diverse roles of DNA methylation in mammalian development and disease. Nat. Rev. Mol. Cell Biol. 20, 590–607. https://doi.org/10.1038/s41580-019-0159-6

GTEx Consortium, Laboratory, Data Analysis &Coordinating Center (LDACC)—Analysis Working Group, Statistical Methods groups—Analysis Working Group, Enhancing GTEx (eGTEx) groups, NIH Common Fund, NIH/NCI, NIH/NHGRI, NIH/NIMH, NIH/NIDA, Biospecimen Collection Source Site—NDRI, Biospecimen Collection Source Site—RPCI, Biospecimen Core Resource—VARI, Brain Bank Repository—University of Miami Brain Endowment Bank, Leidos Biomedical—Project Management, ELSI Study, Genome Browser Data Integration &Visualization—EBI, Genome Browser Data Integration &Visualization—UCSC Genomics Institute, University of California Santa Cruz, Lead analysts:, Laboratory, Data Analysis &Coordinating Center (LDACC):, NIH program management:, Biospecimen collection:, Pathology:, eQTL manuscript working group:, Battle, A., Brown, C.D., Engelhardt, B.E., Montgomery, S.B., 2017. Genetic effects on gene expression across human tissues. Nature 550, 204–213. https://doi.org/10.1038/nature24277

Heintzman, N.D., Stuart, R.K., Hon, G., Fu, Y., Ching, C.W., Hawkins, R.D., Barrera, L.O., Van Calcar, S., Qu, C., Ching, K.A., Wang, W., Weng, Z., Green, R.D., Crawford, G.E., Ren, B., 2007. Distinct and predictive chromatin signatures of transcriptional promoters and enhancers in the human genome. Nat. Genet. 39, 311–318. https://doi.org/10.1038/ng1966

Herbst, A., Jurinovic, V., Krebs, S., Thieme, S.E., Blum, H., Göke, B., Kolligs, F.T., 2014. Comprehensive analysis of β-catenin target genes in colorectal carcinoma cell lines with deregulated Wnt/β-catenin signaling. BMC Genomics 15, 74. https://doi.org/10.1186/1471-2164-15-74

Hoffmann, W., 2015. Current Status on Stem Cells and Cancers of the Gastric Epithelium. Int J Mol Sci 16, 19153–19169. https://doi.org/10.3390/ijms160819153

Holland, A.M., Garcia, S., Naselli, G., Macdonald, R.J., Harrison, L.C., 2013. The Parahox gene Pdx1 is required to maintain positional identity in the adult foregut. Int. J. Dev. Biol. 57, 391–398. https://doi.org/10.1387/ijdb.120048ah

Huang, F.-Y., Chan, A.O.-O., Rashid, A., Wong, D.K.-H., Cho, C.-H., Yuen, M.-F., 2012. Helicobacter pylori induces promoter methylation of E-cadherin via interleukin-1β activation of nitric oxide production in gastric cancer cells. Cancer 118, 4969–4980. https://doi.org/10.1002/cncr.27519

Huang, K.K., Ramnarayanan, K., Zhu, F., Srivastava, S., Xu, C., Tan, A.L.K., Lee, M., Tay, S., Das, K., Xing, M., Fatehullah, A., Alkaff, S.M.F., Lim, T.K.H., Lee, J., Ho, K.Y., Rozen, S.G., Teh, B.T., Barker, N., Chia, C.K., Khor, C., Ooi, C.J., Fock, K.M., So, J., Lim, W.C., Ling, K.L., Ang, T.L., Wong, A., Rao, J., Rajnakova, A., Lim, L.G., Yap, W.M., Teh, M., Yeoh, K.G., Tan, P., 2018. Genomic and Epigenomic Profiling of High-Risk Intestinal Metaplasia Reveals Molecular Determinants of Progression to Gastric Cancer. Cancer Cell 33, 137–150.e5. https://doi.org/10.1016/j.ccell.2017.11.018

Hughes, L.A.E., Melotte, V., de Schrijver, J., de Maat, M., Smit, V.T.H.B.M., Bovée, J.V.M.G., French, P.J., van den Brandt, P.A., Schouten, L.J., de Meyer, T., van Criekinge, W., Ahuja, N., Herman, J.G., Weijenberg, M.P., van Engeland, M., 2013. The CpG island methylator phenotype: what’s in a name? Cancer Res. 73, 5858–5868. https://doi.org/10.1158/0008-5472.CAN-12-4306

Kent, W.J., Sugnet, C.W., Furey, T.S., Roskin, K.M., Pringle, T.H., Zahler, A.M., Haussler, D., 2002. The human genome browser at UCSC. Genome Res. 12, 996–1006. https://doi.org/10.1101/gr.229102

Khan, A., Fornes, O., Stigliani, A., Gheorghe, M., Castro-Mondragon, J.A., van der Lee, R., Bessy, A., Chèneby, J., Kulkarni, S.R., Tan, G., Baranasic, D., Arenillas, D.J., Sandelin, A., Vandepoele, K., Lenhard, B., Ballester, B., Wasserman, W.W., Parcy, F., Mathelier, A., 2018. JASPAR 2018: update of the open-access database of transcription factor binding profiles and its web framework. Nucleic Acids Res. 46, D1284. https://doi.org/10.1093/nar/gkx1188

Kim, T.-H., Shivdasani, R.A., 2016. Stomach development, stem cells and disease. Development 143, 554–565. https://doi.org/10.1242/dev.124891

Krause, L., Nones, K., Loffler, K.A., Nancarrow, D., Oey, H., Tang, Y.H., Wayte, N.J., Patch, A.M., Patel, K., Brosda, S., Manning, S., Lampe, G., Clouston, A., Thomas, J., Stoye, J., Hussey, D.J., Watson, D.I., Lord, R.V., Phillips, W.A., Gotley, D., Smithers, B.M., Whiteman, D.C., Hayward, N.K., Grimmond, S.M., Waddell, N., Barbour, A.P., 2016. Identification of the CIMP-like subtype and aberrant methylation of members of the chromosomal segregation and spindle assembly pathways in esophageal adenocarcinoma. Carcinogenesis 37, 356–365. https://doi.org/10.1093/carcin/bgw018

Larsson, L.I., Madsen, O.D., Serup, P., Jonsson, J., Edlund, H., 1996. Pancreatic-duodenal homeobox 1 -role in gastric endocrine patterning. Mech Dev 60, 175–184. https://doi.org/10.1016/s0925-4773(96)00609-0

Laugesen, A., Højfeldt, J.W., Helin, K., 2019. Molecular Mechanisms Directing PRC2 Recruitment and H3K27 Methylation. Mol. Cell 74, 8–18. https://doi.org/10.1016/j.molcel.2019.03.011

Lieberman-Aiden, E., van Berkum, N.L., Williams, L., Imakaev, M., Ragoczy, T., Telling, A., Amit, I., Lajoie, B.R., Sabo, P.J., Dorschner, M.O., Sandstrom, R., Bernstein, B., Bender, M.A., Groudine, M., Gnirke, A., Stamatoyannopoulos, J., Mirny, L.A., Lander, E.S., Dekker, J., 2009. Comprehensive mapping of long-range interactions reveals folding principles of the human genome. Science 326, 289–293. https://doi.org/10.1126/science.1181369

Lizio, M., Harshbarger, J., Shimoji, H., Severin, J., Kasukawa, T., Sahin, S., Abugessaisa, I., Fukuda, S., Hori, F., Ishikawa-Kato, S., Mungall, C.J., Arner, E., Baillie, J.K., Bertin, N., Bono, H., de Hoon, M., Diehl, A.D., Dimont, E., Freeman, T.C., Fujieda, K., Hide, W., Kaliyaperumal, R., Katayama, T., Lassmann, T., Meehan, T.F., Nishikata, K., Ono, H., Rehli, M., Sandelin, A., Schultes, E.A., ’t Hoen, P.A.C., Tatum, Z., Thompson, M., Toyoda, T., Wright, D.W., Daub, C.O., Itoh, M., Carninci, P., Hayashizaki, Y., Forrest, A.R.R., Kawaji, H., FANTOM consortium, 2015. Gateways to the FANTOM5 promoter level mammalian expression atlas. Genome Biol 16, 22. https://doi.org/10.1186/s13059-014-0560-6

Marttila, S., Kananen, L., Häyrynen, S., Jylhävä, J., Nevalainen, T., Hervonen, A., Jylhä, M., Nykter, M., Hurme, M., 2015. Ageing-associated changes in the human DNA methylome: genomic locations and effects on gene expression. BMC Genomics 16, 179. https://doi.org/10.1186/s12864-015-1381-z

Matsuo, J., Kimura, S., Yamamura, A., Koh, C.P., Hossain, M.Z., Heng, D.L., Kohu, K., Voon, D.C.-C., Hiai, H., Unno, M., So, J.B.Y., Zhu, F., Srivastava, S., Teh, M., Yeoh, K.G., Osato, M., Ito, Y., 2017. Identification of Stem Cells in the Epithelium of the Stomach Corpus and Antrum of Mice. Gastroenterology 152, 218–231.e14. https://doi.org/10.1053/j.gastro.2016.09.018

McCracken, K.W., Aihara, E., Martin, B., Crawford, C.M., Broda, T., Treguier, J., Zhang, X., Shannon, J.M., Montrose, M.H., Wells, J.M., 2017. Wnt/β-catenin promotes gastric fundus specification in mice and humans. Nature 541, 182–187. https://doi.org/10.1038/nature21021

McCracken, K.W., Catá, E.M., Crawford, C.M., Sinagoga, K.L., Schumacher, M., Rockich, B.E., Tsai, Y.-H., Mayhew, C.N., Spence, J.R., Zavros, Y., Wells, J.M., 2014. Modelling human development and disease in pluripotent stem-cell-derived gastric organoids. Nature 516, 400–404. https://doi.org/10.1038/nature13863

Monk, D., 2015. Germline-derived DNA methylation and early embryo epigenetic reprogramming: The selected survival of imprints. Int. J. Biochem. Cell Biol. 67, 128–138. https://doi.org/10.1016/j.biocel.2015.04.014

Morak, M., Schackert, H.K., Rahner, N., Betz, B., Ebert, M., Walldorf, C., Royer-Pokora, B., Schulmann, K., von Knebel-Doeberitz, M., Dietmaier, W., Keller, G., Kerker, B., Leitner, G., Holinski-Feder, E., 2008. Further evidence for heritability of an epimutation in one of 12 cases with MLH1 promoter methylation in blood cells clinically displaying HNPCC. Eur. J. Hum. Genet. 16, 804–811. https://doi.org/10.1038/ejhg.2008.25

Moran, S., Arribas, C., Esteller, M., 2016. Validation of a DNA methylation microarray for 850,000 CpG sites of the human genome enriched in enhancer sequences. Epigenomics 8, 389–399. https://doi.org/10.2217/epi.15.114

Morris, T.J., Butcher, L.M., Feber, A., Teschendorff, A.E., Chakravarthy, A.R., Wojdacz, T.K., Beck, S., 2014. ChAMP: 450k Chip Analysis Methylation Pipeline. Bioinformatics 30, 428–430. https://doi.org/10.1093/bioinformatics/btt684

Nanki, K., Toshimitsu, K., Takano, A., Fujii, M., Shimokawa, M., Ohta, Y., Matano, M., Seino, T., Nishikori, S., Ishikawa, K., Kawasaki, K., Togasaki, K., Takahashi, S., Sukawa, Y., Ishida, H., Sugimoto, S., Kawakubo, H., Kim, J., Kitagawa, Y., Sekine, S., Koo, B.-K., Kanai, T., Sato, T., 2018. Divergent Routes toward Wnt and R-spondin Niche Independency during Human Gastric Carcinogenesis. Cell 174, 856–869.e17. https://doi.org/10.1016/j.cell.2018.07.027

Ng, R.K., Dean, W., Dawson, C., Lucifero, D., Madeja, Z., Reik, W., Hemberger, M., 2008. Epigenetic restriction of embryonic cell lineage fate by methylation of Elf5. Nat. Cell Biol. 10, 1280–1290. https://doi.org/10.1038/ncb1786

Niwa, T., Tsukamoto, T., Toyoda, T., Mori, A., Tanaka, H., Maekita, T., Ichinose, M., Tatematsu, M., Ushijima, T., 2010. Inflammatory processes triggered by Helicobacter pylori infection cause aberrant DNA methylation in gastric epithelial cells. Cancer Res. 70, 1430–1440. https://doi.org/10.1158/0008-5472.CAN-09-2755

Nookaew, I., Thorell, K., Worah, K., Wang, S., Hibberd, M.L., Sjövall, H., Pettersson, S., Nielsen, J., Lundin, S.B., 2013. Transcriptome signatures in Helicobacter pylori-infected mucosa identifies acidic mammalian chitinase loss as a corpus atrophy marker. BMC Med Genomics 6, 41. https://doi.org/10.1186/1755-8794-6-41

Offield, M.F., Jetton, T.L., Labosky, P.A., Ray, M., Stein, R.W., Magnuson, M.A., Hogan, B.L., Wright, C.V., 1996. PDX-1 is required for pancreatic outgrowth and differentiation of the rostral duodenum. Development 122, 983–995.

Oishi, I., Suzuki, H., Onishi, N., Takada, R., Kani, S., Ohkawara, B., Koshida, I., Suzuki, K., Yamada, G., Schwabe, G.C., Mundlos, S., Shibuya, H., Takada, S., Minami, Y., 2003. The receptor tyrosine kinase Ror2 is involved in non-canonical Wnt5a/JNK signalling pathway. Genes to Cells 8, 645–654. https://doi.org/10.1046/j.1365-2443.2003.00662.x

Pekowska, A., Benoukraf, T., Zacarias-Cabeza, J., Belhocine, M., Koch, F., Holota, H., Imbert, J., Andrau, J.-C., Ferrier, P., Spicuglia, S., 2011. H3K4 tri-methylation provides an epigenetic signature of active enhancers. EMBO J 30, 4198–4210. https://doi.org/10.1038/emboj.2011.295

Pereira, B., Oliveira, C., David, L., Almeida, R., 2009. CDX2 promoter methylation is not associated with mRNA expression. Int. J. Cancer 125, 1739–1742. https://doi.org/10.1002/ijc.24544

R Core Team (2017), n.d. R: A language and environment for statistical computing. R Foundation for Statistical Computing, Vienna, Austria.

Rada-Iglesias, A., Bajpai, R., Swigut, T., Brugmann, S.A., Flynn, R.A., Wysocka, J., 2011. A unique chromatin signature uncovers early developmental enhancers in humans. Nature 470, 279–283. https://doi.org/10.1038/nature09692

Rajasekhar, V.K., Begemann, M., 2007. Concise review: roles of polycomb group proteins in development and disease: a stem cell perspective. Stem Cells 25, 2498–2510. https://doi.org/10.1634/stemcells.2006-0608

Ramsey, V.G., Doherty, J.M., Chen, C.C., Stappenbeck, T.S., Konieczny, S.F., Mills, J.C., 2007. The maturation of mucus-secreting gastric epithelial progenitors into digestive-enzyme secreting zymogenic cells requires Mist1. Development 134, 211–222. https://doi.org/10.1242/dev.02700

Ritchie, M.E., Phipson, B., Wu, D., Hu, Y., Law, C.W., Shi, W., Smyth, G.K., 2015. limma powers differential expression analyses for RNA-sequencing and microarray studies. Nucleic Acids Res. 43, e47. https://doi.org/10.1093/nar/gkv007

Ritchie, M.E., Silver, J., Oshlack, A., Holmes, M., Diyagama, D., Holloway, A., Smyth, G.K., 2007. A comparison of background correction methods for two-colour microarrays. Bioinformatics 23, 2700– 2707. https://doi.org/10.1093/bioinformatics/btm412

Roadmap Epigenomics Consortium, Kundaje, A., Meuleman, W., Ernst, J., Bilenky, M., Yen, A., Heravi-Moussavi, A., Kheradpour, P., Zhang, Z., Wang, J., Ziller, M.J., Amin, V., Whitaker, J.W., Schultz, M.D., Ward, L.D., Sarkar, A., Quon, G., Sandstrom, R.S., Eaton, M.L., Wu, Y.-C., Pfenning, A.R., Wang, X., Claussnitzer, M., Liu, Y., Coarfa, C., Harris, R.A., Shoresh, N., Epstein, C.B., Gjoneska, E., Leung, D., Xie, W., Hawkins, R.D., Lister, R., Hong, C., Gascard, P., Mungall, A.J., Moore, R., Chuah, E., Tam, A., Canfield, T.K., Hansen, R.S., Kaul, R., Sabo, P.J., Bansal, M.S., Carles, A., Dixon, J.R., Farh, K.-H., Feizi, S., Karlic, R., Kim, A.-R., Kulkarni, A., Li, D., Lowdon, R., Elliott, G., Mercer, T.R., Neph, S.J., Onuchic, V., Polak, P., Rajagopal, N., Ray, P., Sallari, R.C., Siebenthall, K.T., Sinnott-Armstrong, N.A., Stevens, M., Thurman, R.E., Wu, J., Zhang, B., Zhou, X., Beaudet, A.E., Boyer, L.A., De Jager, P.L., Farnham, P.J., Fisher, S.J., Haussler, D., Jones, S.J.M., Li, W., Marra, M.A., McManus, M.T., Sunyaev, S., Thomson, J.A., Tlsty, T.D., Tsai, L.-H., Wang, W., Waterland, R.A., Zhang, M.Q., Chadwick, L.H., Bernstein, B.E., Costello, J.F., Ecker, J.R., Hirst, M., Meissner, A., Milosavljevic, A., Ren, B., Stamatoyannopoulos, J.A., Wang, T., Kellis, M., 2015. Integrative analysis of 111 reference human epigenomes. Nature 518, 317–330. https://doi.org/10.1038/nature14248

Saldanha, J., Singh, J., Mahadevan, D., 1998. Identification of a frizzled-like cysteine rich domain in the extracellular region of developmental receptor tyrosine kinases. Protein Science 7, 1632–1635. https://doi.org/10.1002/pro.5560070718

Schlaermann, P., Toelle, B., Berger, H., Schmidt, S.C., Glanemann, M., Ordemann, J., Bartfeld, S., Mollenkopf, H.J., Meyer, T.F., 2016. A novel human gastric primary cell culture system for modelling Helicobacter pylori infection in vitro. Gut 65, 202–213. https://doi.org/10.1136/gutjnl-2014-307949

Schlesinger, Y., Straussman, R., Keshet, I., Farkash, S., Hecht, M., Zimmerman, J., Eden, E., Yakhini, Z., Ben-Shushan, E., Reubinoff, B.E., Bergman, Y., Simon, I., Cedar, H., 2007. Polycomb-mediated methylation on Lys27 of histone H3 pre-marks genes for de novo methylation in cancer. Nat. Genet. 39, 232–236. https://doi.org/10.1038/ng1950

Schulte, D., Geerts, D., 2019. MEIS transcription factors in development and disease. Development 146. https://doi.org/10.1242/dev.174706

Sheaffer, K.L., Kim, R., Aoki, R., Elliott, E.N., Schug, J., Burger, L., Schübeler, D., Kaestner, K.H., 2014. DNA methylation is required for the control of stem cell differentiation in the small intestine. Genes Dev. 28, 652–664. https://doi.org/10.1101/gad.230318.113

Sheffield, N.C., Bock, C., 2016. LOLA: enrichment analysis for genomic region sets and regulatory elements in R and Bioconductor. Bioinformatics 32, 587–589. https://doi.org/10.1093/bioinformatics/btv612

Siggens, L., Ekwall, K., 2014. Epigenetics, chromatin and genome organization: recent advances from the ENCODE project. J. Intern. Med. 276, 201–214. https://doi.org/10.1111/joim.12231

Smith, Z.D., Shi, J., Gu, H., Donaghey, J., Clement, K., Cacchiarelli, D., Gnirke, A., Michor, F., Meissner, A., 2017. Epigenetic restriction of extra-embryonic lineages mirrors the somatic transition to cancer. Nature 549, 543–547. https://doi.org/10.1038/nature23891

Stange, D.E., Koo, B.-K., Huch, M., Sibbel, G., Basak, O., Lyubimova, A., Kujala, P., Bartfeld, S., Koster, J., Geahlen, J.H., Peters, P.J., van Es, J.H., van de Wetering, M., Mills, J.C., Clevers, H., 2013. Differentiated Troy+ chief cells act as reserve stem cells to generate all lineages of the stomach epithelium. Cell 155, 357–368. https://doi.org/10.1016/j.cell.2013.09.008

Stoffers, D.A., Heller, R.S., Miller, C.P., Habener, J.F., 1999. Developmental Expression of the Homeodomain Protein IDX-1 in Mice Transgenic for an IDX-1 Promoter/lacZ Transcriptional Reporter. Endocrinology 140, 5374–5381. https://doi.org/10.1210/endo.140.11.7122

Stringer, E.J., Duluc, I., Saandi, T., Davidson, I., Bialecka, M., Sato, T., Barker, N., Clevers, H., Pritchard, C.A., Winton, D.J., Wright, N.A., Freund, J.-N., Deschamps, J., Beck, F., 2012. Cdx2 determines the fate of postnatal intestinal endoderm. Development 139, 465–474. https://doi.org/10.1242/dev.070722

Subramanian, A., Tamayo, P., Mootha, V.K., Mukherjee, S., Ebert, B.L., Gillette, M.A., Paulovich, A., Pomeroy, S.L., Golub, T.R., Lander, E.S., Mesirov, J.P., 2005. Gene set enrichment analysis: a knowledge-based approach for interpreting genome-wide expression profiles. Proc. Natl. Acad. Sci. U.S.A. 102, 15545–15550. https://doi.org/10.1073/pnas.0506580102

Szklarczyk, D., Franceschini, A., Wyder, S., Forslund, K., Heller, D., Huerta-Cepas, J., Simonovic, M., Roth, A., Santos, A., Tsafou, K.P., Kuhn, M., Bork, P., Jensen, L.J., von Mering, C., 2015. STRING v10: protein-protein interaction networks, integrated over the tree of life. Nucleic Acids Res. 43, D447–452. https://doi.org/10.1093/nar/gku1003

Tian, Y., Morris, T.J., Webster, A.P., Yang, Z., Beck, S., Feber, A., Teschendorff, A.E., 2017. ChAMP: updated methylation analysis pipeline for Illumina BeadChips. Bioinformatics 33, 3982–3984. https://doi.org/10.1093/bioinformatics/btx513

Uemura, N., Okamoto, S., Yamamoto, S., Matsumura, N., Yamaguchi, S., Yamakido, M., Taniyama, K., Sasaki, N., Schlemper, R.J., 2001. Helicobacter pylori infection and the development of gastric cancer. N. Engl. J. Med. 345, 784–789. https://doi.org/10.1056/NEJMoa001999

Uhlén, M., Fagerberg, L., Hallström, B.M., Lindskog, C., Oksvold, P., Mardinoglu, A., Sivertsson, Å., Kampf, C., Sjöstedt, E., Asplund, A., Olsson, I., Edlund, K., Lundberg, E., Navani, S., Szigyarto, C.A.-K., Odeberg, J., Djureinovic, D., Takanen, J.O., Hober, S., Alm, T., Edqvist, P.-H., Berling, H., Tegel, H., Mulder, J., Rockberg, J., Nilsson, P., Schwenk, J.M., Hamsten, M., von Feilitzen, K., Forsberg, M., Persson, L., Johansson, F., Zwahlen, M., von Heijne, G., Nielsen, J., Pontén, F., 2015. Proteomics. Tissue-based map of the human proteome. Science 347, 1260419. https://doi.org/10.1126/science.1260419

Vaquerizas, J.M., Kummerfeld, S.K., Teichmann, S.A., Luscombe, N.M., 2009. A census of human transcription factors: function, expression and evolution. Nat. Rev. Genet. 10, 252–263. https://doi.org/10.1038/nrg2538

Vivian, J., Rao, A.A., Nothaft, F.A., Ketchum, C., Armstrong, J., Novak, A., Pfeil, J., Narkizian, J., Deran, A.D., Musselman-Brown, A., Schmidt, H., Amstutz, P., Craft, B., Goldman, M., Rosenbloom, K., Cline, M., O’Connor, B., Hanna, M., Birger, C., Kent, W.J., Patterson, D.A., Joseph, A.D., Zhu, J., Zaranek, S., Getz, G., Haussler, D., Paten, B., 2017. Toil enables reproducible, open source, big biomedical data analyses. Nat Biotechnol 35, 314–316. https://doi.org/10.1038/nbt.3772

Wölffling, S., Daddi, A.A., Imai-Matsushima, A., Fritsche, K., Goosmann, C., Traulsen, J., Lisle, R., Schmid, M., Reines-Benassar, M.D.M., Pfannkuch, L., Brinkmann, V., Bornschein, J., Malfertheiner, P., Ordemann, J., Link, A., Meyer, T.F., Boccellato, F., 2021. EGF and BMPs Govern Differentiation and Patterning in Human Gastric Glands. Gastroenterology. https://doi.org/10.1053/j.gastro.2021.04.062

Woo, H.D., Fernandez-Jimenez, N., Ghantous, A., Degli Esposti, D., Cuenin, C., Cahais, V., Choi, I.J., Kim, Y.-I., Kim, J., Herceg, Z., 2018. Genome-wide profiling of normal gastric mucosa identifies Helicobacter pylori- and cancer-associated DNA methylome changes. Int. J. Cancer 143, 597–609. https://doi.org/10.1002/ijc.31381

Yamashita, S., Nanjo, S., Rehnberg, E., Iida, N., Takeshima, H., Ando, T., Maekita, T., Sugiyama, T., Ushijima, T., 2019. Distinct DNA methylation targets by aging and chronic inflammation: a pilot study using gastric mucosa infected with Helicobacter pylori. Clin Epigenetics 11, 191. https://doi.org/10.1186/s13148-019-0789-8

Yoda, Y., Takeshima, H., Niwa, T., Kim, J.G., Ando, T., Kushima, R., Sugiyama, T., Katai, H., Noshiro, H., Ushijima, T., 2015. Integrated analysis of cancer-related pathways affected by genetic and epigenetic alterations in gastric cancer. Gastric Cancer 18, 65–76. https://doi.org/10.1007/s10120-014-0348-0

Zhou, W., Dinh, H.Q., Ramjan, Z., Weisenberger, D.J., Nicolet, C.M., Shen, H., Laird, P.W., Berman, B.P., 2018. DNA methylation loss in late-replicating domains is linked to mitotic cell division. Nat. Genet. 50, 591–602. https://doi.org/10.1038/s41588-018-0073-4

