## Additional Figures for "DNA methylation in human gastric epithelial cells allows cell type-related plasticity and defines regional identity"

### Suppl. Fig.1

A

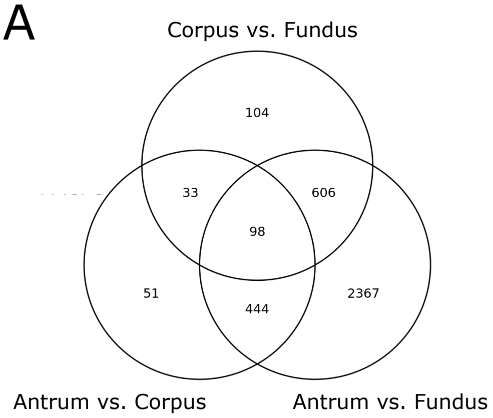


B

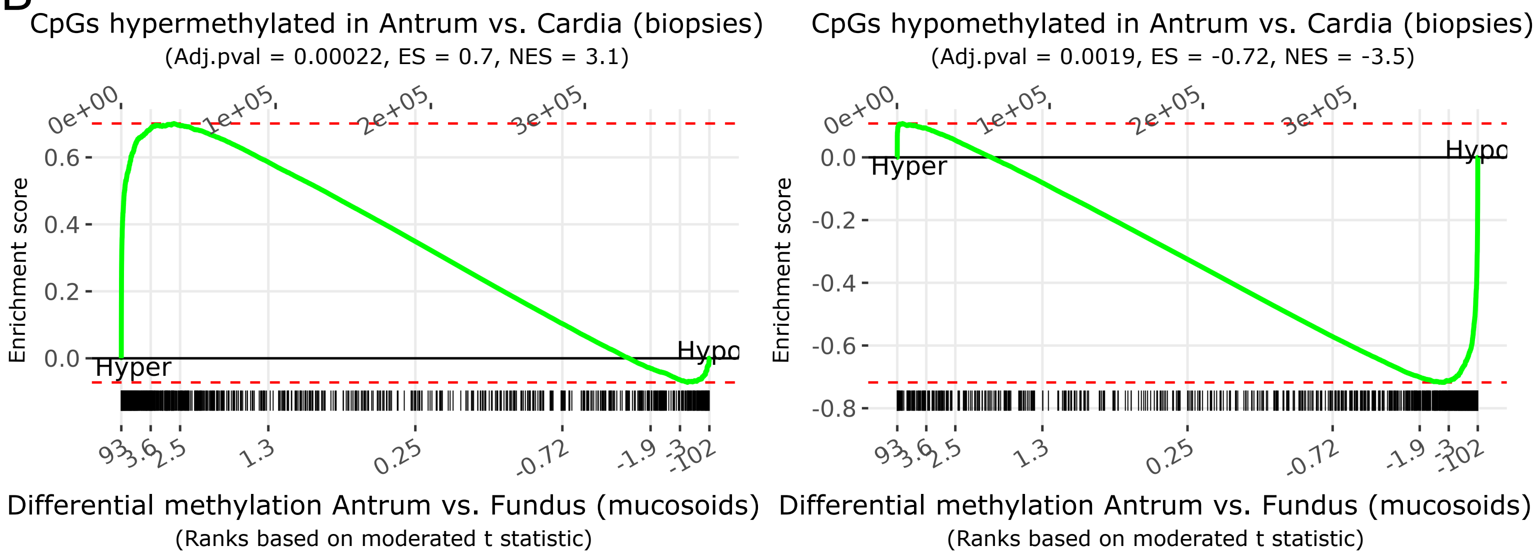


C

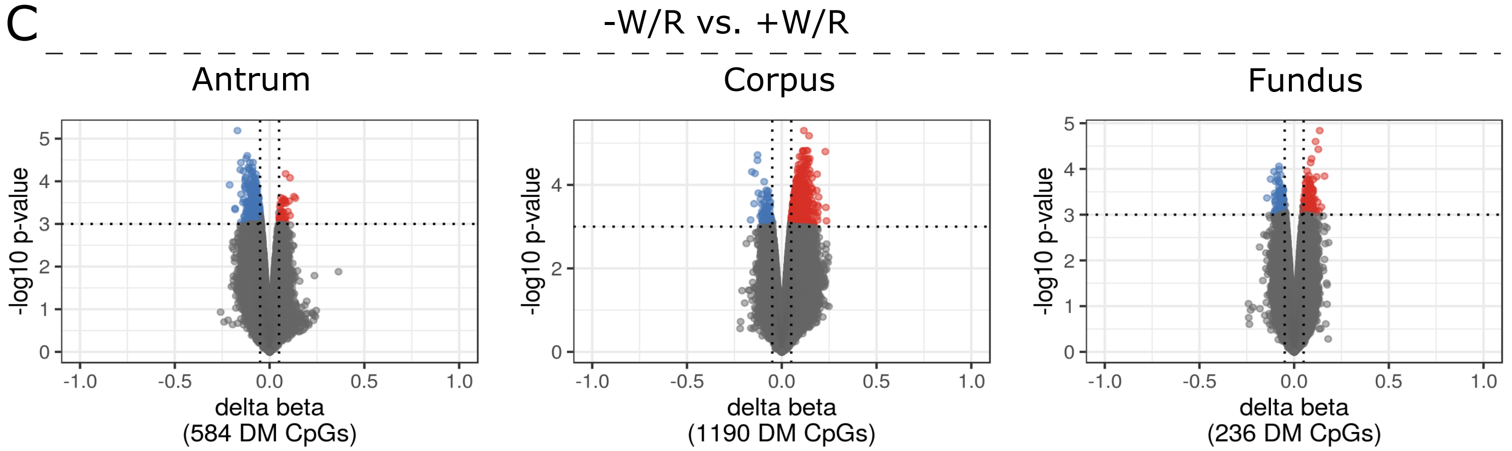


D

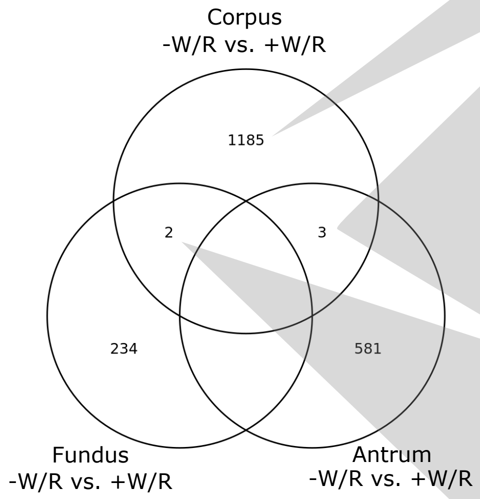


| CG ID (Corpus) | logFC | p-value | FDR | CHR | MAPINFO | gene | feature | cgi |
|---|---|---|---|---|---|---|---|---|
| cg10758260 | 0.1 | 0.0005 | 0.3645 | 13 | 24242176 | TNFRSF19 | Body | opensea |

| CG ID (Antrum) | logFC | p-value | FDR | CHR | MAPINFO | gene | feature | cgi |
|---|---|---|---|---|---|---|---|---|
| cg26174018 | -0.09 | 0.0004 | 0.7266 | 14 | 38876825 | | IGR | opensea |
| cg05089925 | -0.12 | 0.0005 | 0.7266 | 10 | 4477161 | | IGR | opensea |
| cg19843226 | -0.13 | 0.0008 | 0.7266 | 14 | 76836781 | ESRRB | TSS1500 | opensea |

| CG ID (Corpus) | logFC | p-value | FDR | CHR | MAPINFO | gene | feature | cgi |
|---|---|---|---|---|---|---|---|---|
| cg26174018 | 0.07 | 0.0007 | 0.3805 | 14 | 38876825 | | IGR | opensea |
| cg05089925 | 0.11 | 0.0007 | 0.3645 | 10 | 4477161 | | IGR | opensea |
| cg19843226 | 0.1 | 0.001 | 0.3645 | 14 | 76836781 | ESRRB | TSS1500 | opensea |

| CG ID (Corpus) | logFC | p-value | FDR | CHR | MAPINFO | gene | feature | cgi |
|---|---|---|---|---|---|---|---|---|
| cg26613034 | 0.13 | 2.38E-05 | 0.3645 | 6 | 13634262 | RANBP9 | Body | opensea |
| cg08376643 | 0.16 | 0.0009 | 0.3645 | 10 | 64880766 | | IGR | opensea |

| CG ID (Fundus) | logFC | p-value | FDR | CHR | MAPINFO | gene | feature | cgi |
|---|---|---|---|---|---|---|---|---|
| cg26613034 | 0.12 | 0.0009 | 1 | 6 | 13634262 | RANBP9 | Body | opensea |
| cg08376643 | 0.12 | 0.0001 | 1 | 10 | 64880766 | | IGR | opensea |

### Suppl. Fig.2

A

| | Antrum vs. Corpus | Antrum vs. Fundus | Corpus vs. Fundus | unique between all regions | Stomach vs. Esophagus | Stomach vs. Small intestine | Stomach vs. Colon |
|---|---|---|---|---|---|---|---|
| Hypermethylated genes | 20 | 161 | 45 | 170 | 2,859 | 1,618 | 1,906 |
| Hypomethylated genes | 32 | 149 | 30 | 159 | 3,395 | 4,538 | 3,720 |
| Hypermethylated promoters | 6 | 45 | 11 | 48 | 1,071 | 619 | 770 |
| Hypomethylated promoters | 9 | 54 | 17 | 57 | 1,367 | 1,956 | 1,545 |

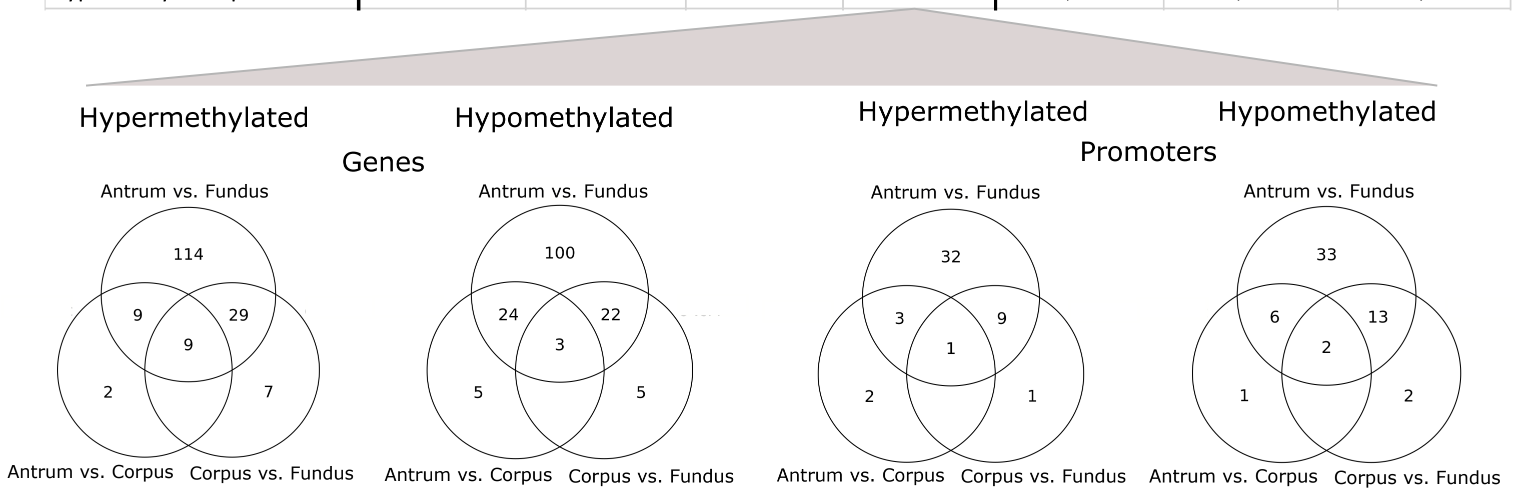


B

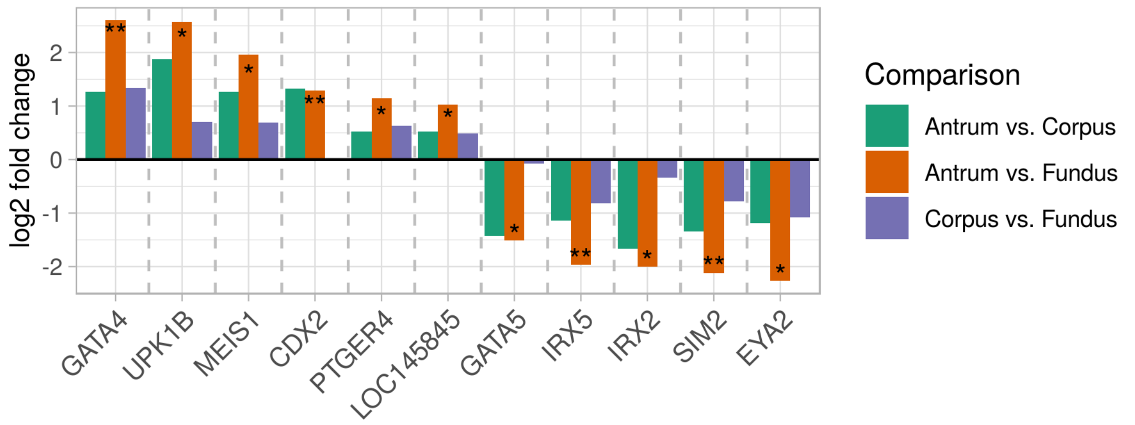


C

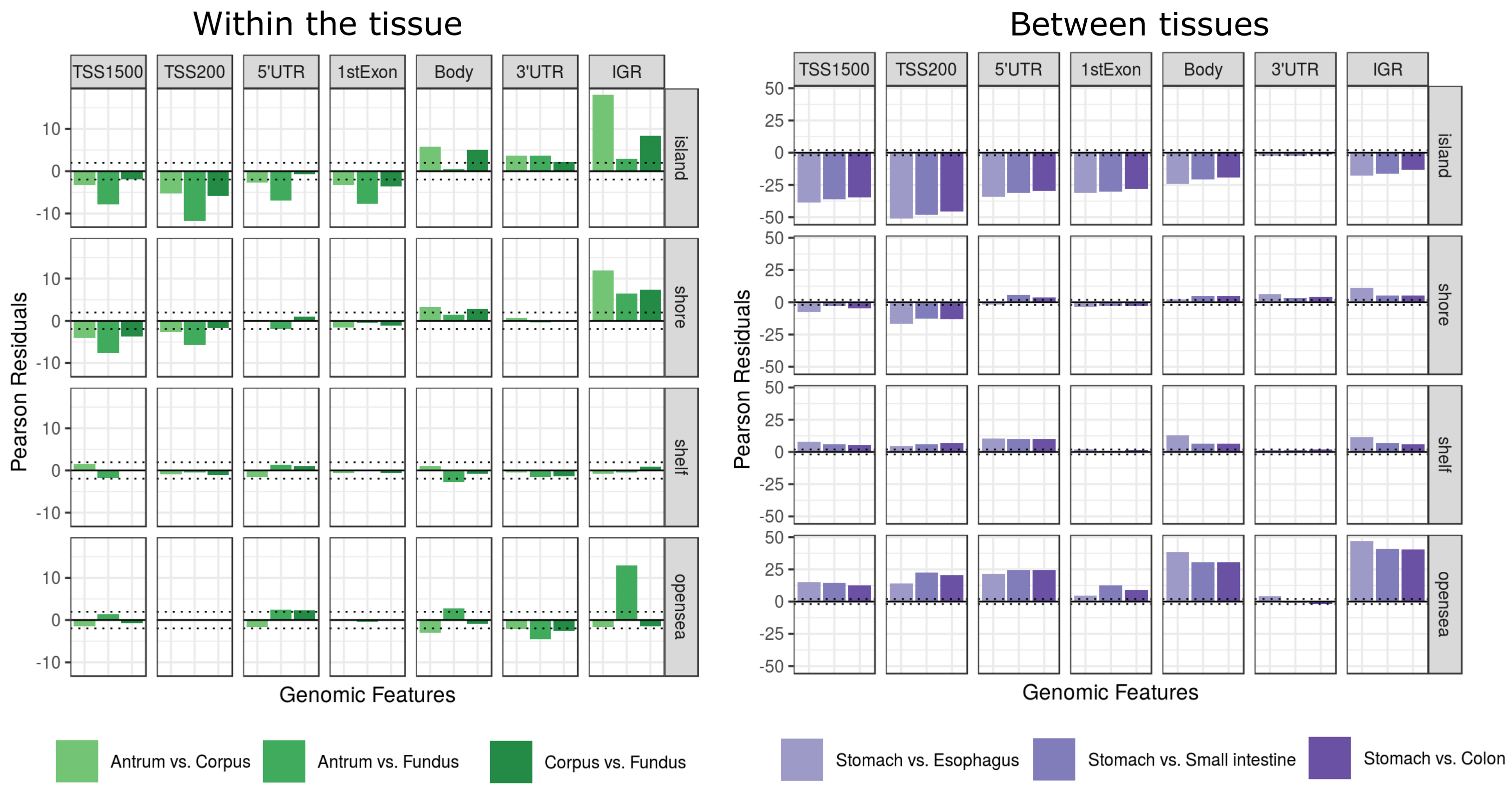

Suppl. Fig. 3

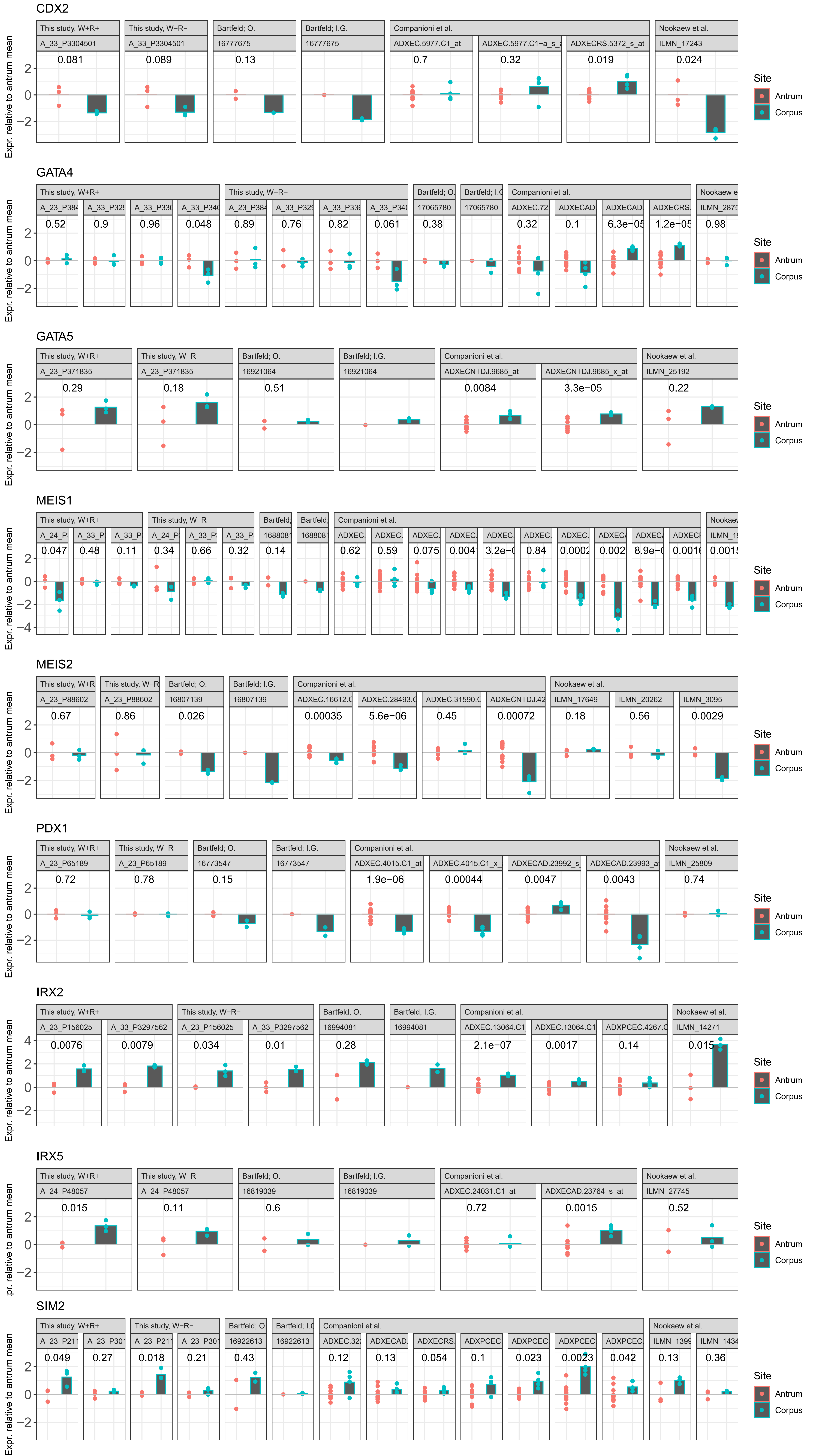
CDX2
GATA4
GATA5
MEIS1
MEIS2
PDX1
IRX2
IRX5
SIM2
Expr. relative to antrum mean
Site
Antrum
Corpus
This study, W+R+
This study, W−R−
Bartfeld; O.
Bartfeld; I.G.
Companioni et al.
Nookaew et al.

### Suppl. Fig.4

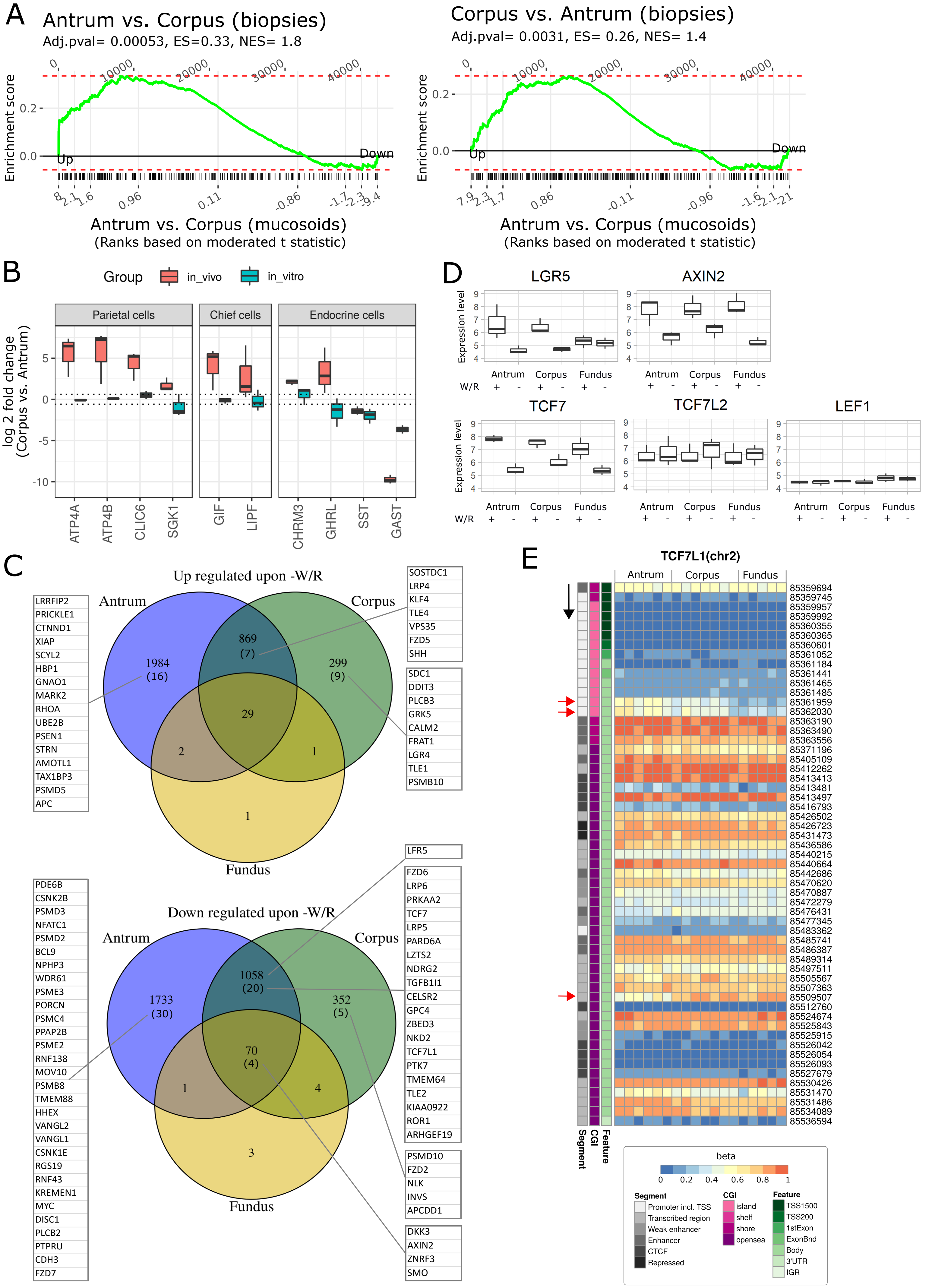

A
Antrum vs. Corpus (biopsies)
Adj.pval= 0.00053, ES=0.33, NES= 1.8
Enrichment score
Up
Down
Antrum vs. Corpus (mucosoids)
(Ranks based on moderated t statistic)
Corpus vs. Antrum (biopsies)
Adj.pval= 0.0031, ES= 0.26, NES= 1.4
Enrichment score
Up
Down
Antrum vs. Corpus (mucosoids)
(Ranks based on moderated t statistic)
B
Group
in_vivo
in_vitro
Parietal cells
Chief cells
Endocrine cells
log 2 fold change
(Corpus vs. Antrum)
ATP4A
ATP4B
CLIC6
SGK1
GIF
LIPF
CHRM3
GHRL
SST
GAST
D
LGR5
AXIN2
TCF7
TCF7L2
LEF1
Expression level
Antrum
Corpus
Fundus
W/R
C
Up regulated upon -W/R
Antrum
Corpus
Fundus
1984 (16)
869 (7)
299 (9)
29
2
1
1
LRRFIP2
PRICKLE1
CTNND1
XIAP
SCYL2
HBP1
GNAO1
MARK2
RHOA
UBE2B
PSEN1
STRN
AMOTL1
TAX1BP3
PSMD5
APC
SOSTDC1
LRP4
KLF4
TLE4
VPS35
FZD5
SHH
SDC1
DDIT3
PLCB3
GRK5
CALM2
FRAT1
LGR4
TLE1
PSMB10
Down regulated upon -W/R
Antrum
Corpus
Fundus
1733 (30)
1058 (20)
352 (5)
70 (4)
1
4
3
PDE6B
CSNK2B
PSMD3
NFATC1
PSMD2
BCL9
NPHP3
WDR61
PSME3
PORCN
PSMC4
PPAP2B
PSME2
RNF138
MOV10
PSMB8
TMEM88
HHEX
VANGL2
VANGL1
CSNK1E
RGS19
RNF43
KREMEN1
MYC
DISC1
PLCB2
PTPRU
CDH3
FZD7
LFR5
FZD6
LRP6
PRKAA2
TCF7
LRP5
PARD6A
LZTS2
NDRG2
TGFB1I1
CELSR2
GPC4
ZBED3
NKD2
TCF7L1
PTK7
TMEM64
TLE2
KIAA0922
ROR1
ARHGEF19
PSMD10
FZD2
NLK
INVS
APCDD1
DKK3
AXIN2
ZNRF3
SMO
E
TCF7L1(chr2)
Antrum
Corpus
Fundus
85359694
85359745
85359957
85359992
85360355
85360365
85360601
85361052
85361184
85361441
85361465
85361485
85361959
85362030
85363190
85363490
85363556
85371196
85405109
85412262
85413413
85413481
85413497
85416793
85426502
85426723
85431473
85436586
85440215
85440664
85442686
85470620
85470887
85472279
85476431
85477345
85483362
85485741
85486387
85489314
85497511
85505567
85507363
85509507
85512760
85524674
85525843
85525915
85526042
85526054
85526093
85527679
85530426
85531470
85531486
85534089
85536594
Segment
CGI
Feature
beta
0
0.2
0.4
0.6
0.8
1
Segment
Promoter incl. TSS
Transcribed region
Weak enhancer
Enhancer
CTCF
Repressed
CGI
island
shelf
shore
opensea
Feature
TSS1500
TSS200
1stExon
ExonBnd
Body
3'UTR
IGR

### Suppl. Fig.5

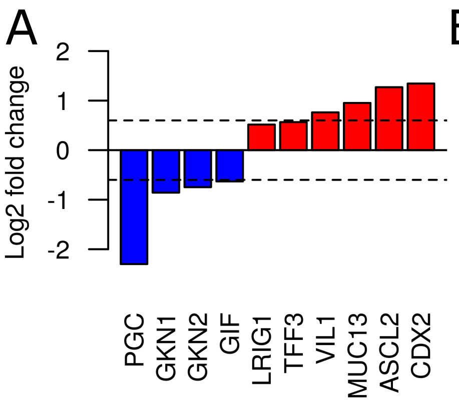


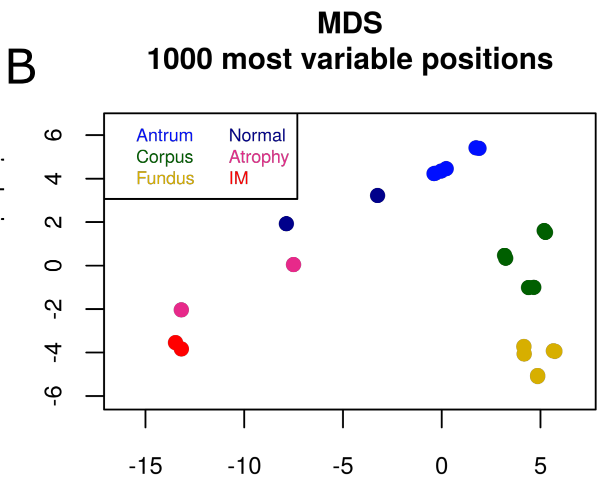


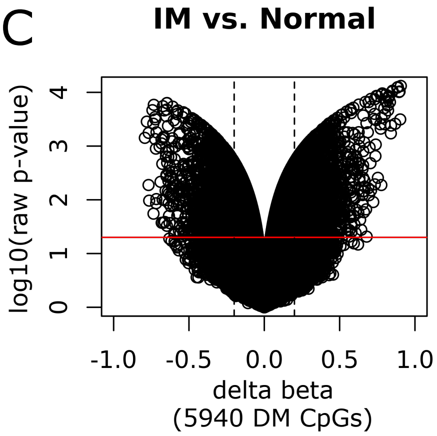


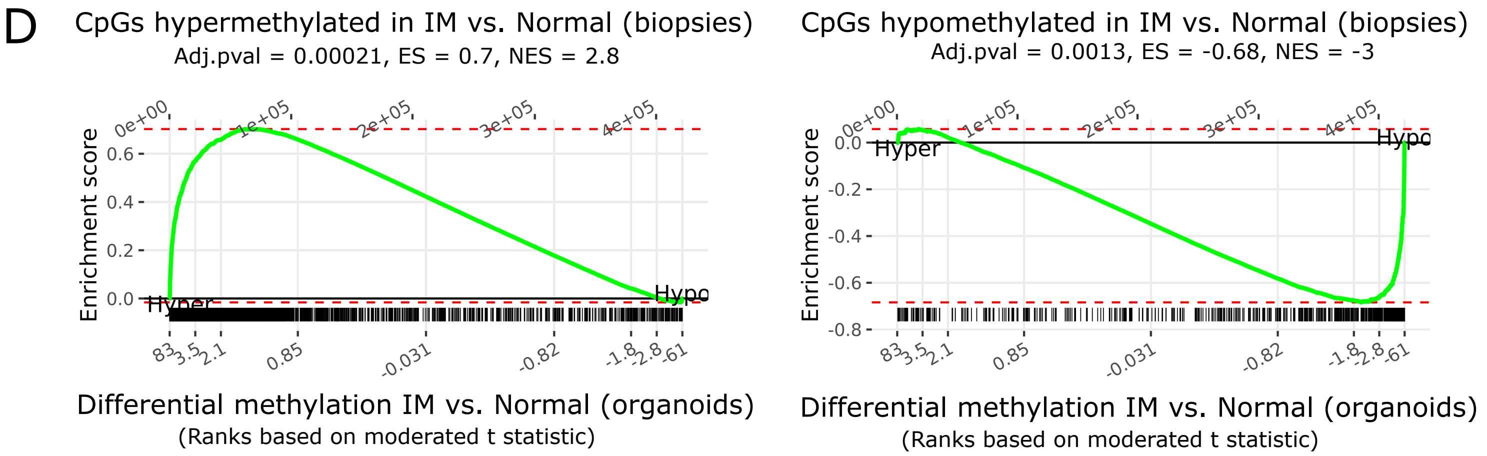


E

**All antral samples of IM**

| | CpGs | | Genes | | Promoters | | Enhancer | |
|---|---|---|---|---|---|---|---|---|
| | Hyper | Hypo | Hyper | Hypo | Hyper | Hypo | Hyper | Hypo |
| ex vivo | 3586 | 2354 | 441 | 215 | 191 | 84 | 118 | 111 |
| in vivo | 14023 | 1199 | 1592 | 94 | 1227 | 37 | 285 | 40 |
| shared | 507 | 107 | 157 | 12 | 74 | 3 | 6 | 1 |
| same CpG | | | 111 | 7 | 64 | 3 | 2 | 1 |

**Antral samples (high methylation cluster) of IM**

| | CpGs | | Genes | | Promoters | | Enhancer | |
|---|---|---|---|---|---|---|---|---|
| | Hyper | Hypo | Hyper | Hypo | Hyper | Hypo | Hyper | Hypo |
| ex vivo | 3586 | 2354 | 441 | 215 | 191 | 84 | 118 | 111 |
| in vivo | 24552 | 5358 | 2274 | 621 | 1748 | 236 | 833 | 235 |
| shared | 767 | 273 | 213 | 61 | 91 | 15 | 24 | 12 |
| same CpG | | | 157 | 38 | 81 | 12 | 7 | 6 |

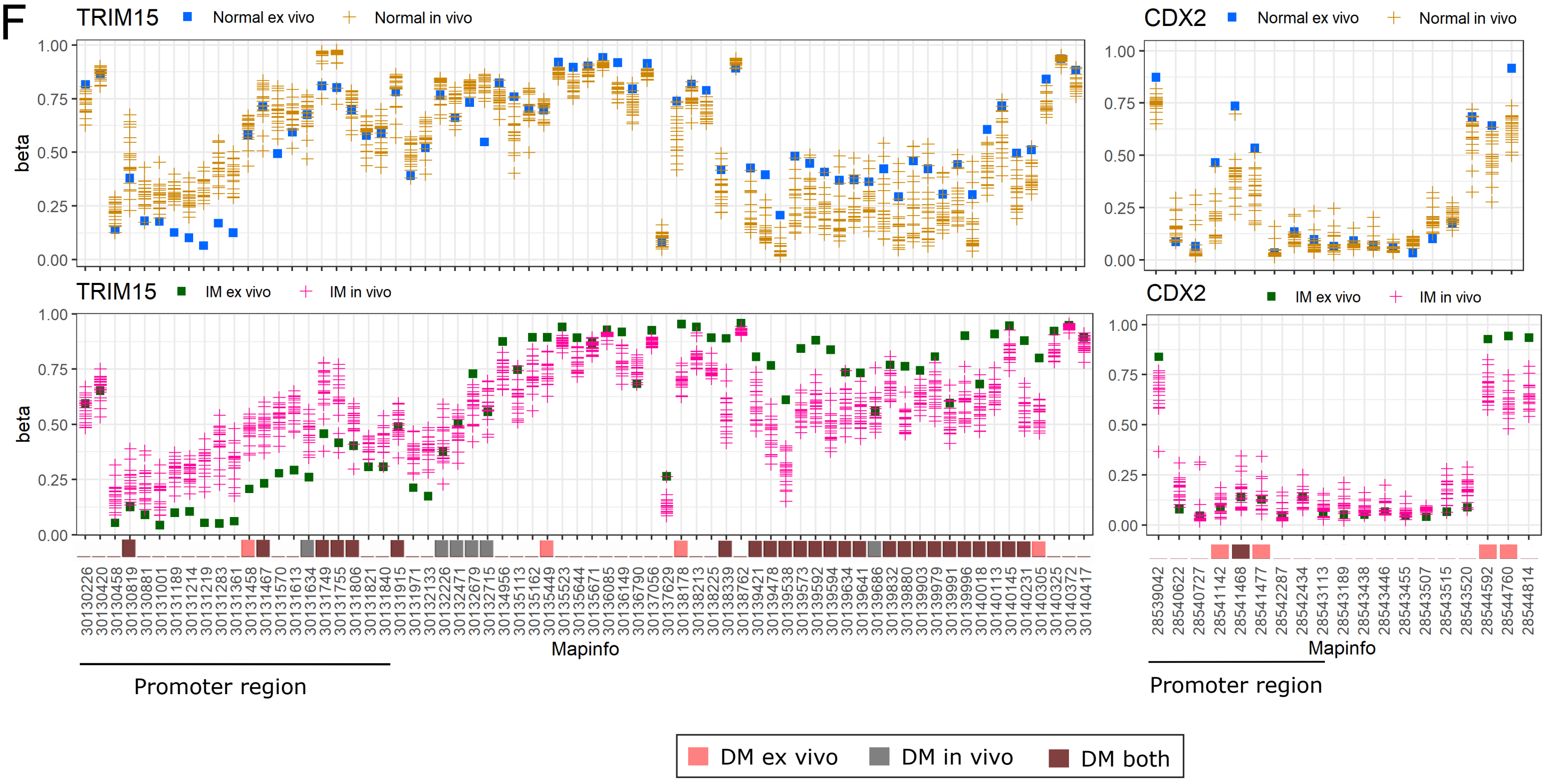

### Suppl. Fig.6

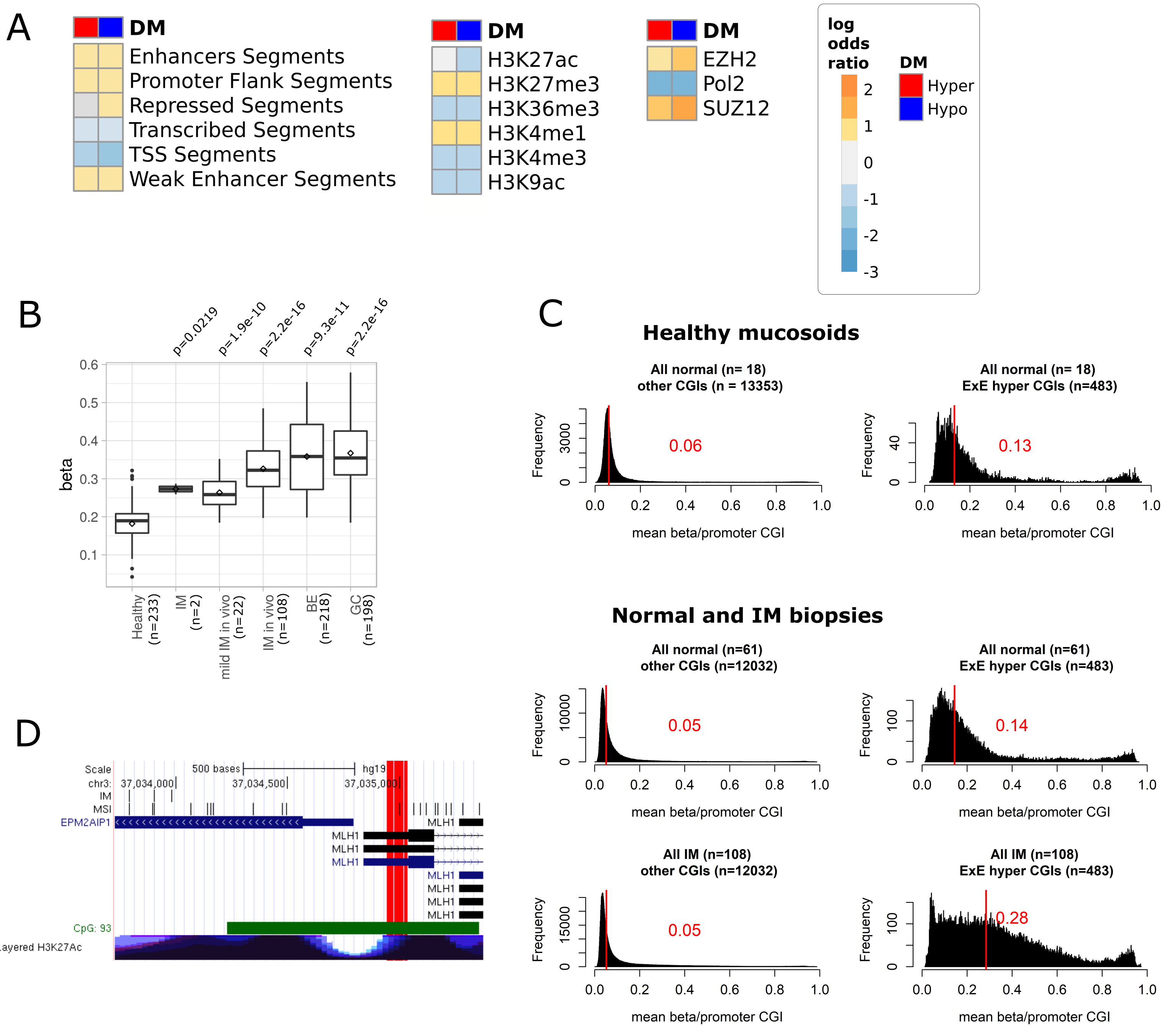
A
DM
Enhancers Segments
Promoter Flank Segments
Repressed Segments
Transcribed Segments
TSS Segments
Weak Enhancer Segments
H3K27ac
H3K27me3
H3K36me3
H3K4me1
H3K4me3
H3K9ac
EZH2
Pol2
SUZ12
log odds ratio
2
1
0
-1
-2
-3
Hyper
Hypo
B
p=0.0219
p=1.9e-10
p=2.2e-16
p=9.3e-11
p=2.2e-16
beta
Healthy (n=233)
IM (n=2)
mild IM in vivo (n=22)
IM in vivo (n=108)
BE (n=218)
GC (n=198)
C
Healthy mucosoids
All normal (n= 18) other CGIs (n = 13353)
0.06
All normal (n= 18) ExE hyper CGIs (n=483)
0.13
Normal and IM biopsies
All normal (n=61) other CGIs (n=12032)
0.05
All normal (n=61) ExE hyper CGIs (n=483)
0.14
All IM (n=108) other CGIs (n=12032)
0.05
All IM (n=108) ExE hyper CGIs (n=483)
0.28
Frequency
mean beta/promoter CGI
D
Scale
500 bases
hg19
chr3:
37,034,000
37,034,500
37,035,000
IM
MSI
EPM2AIP1
MLH1
CpG: 93
Layered H3K27Ac

### Supplementary Figure legends

#### Supplementary Figure 1

**A)** Venn diagram of differentially methylated (DM) CpGs by regional comparisons of the stomach. **B)** Enrichment of sets of CpGs hypermethylated (left) or hypomethylated (right) between antrum and cardia biopsies compared to differential methylation of CpGs between antrum and fundus mucosoids. Moderated t-scores were used for DM CpG ranking. ES - enrichment score; NES - normalized enrichment score. Hyper, Hypo indicate the direction of differential methylation in the antrum vs. fundus mucosoids comparison. **C)** Volcano plots representing comparisons between differentiated (-W/R) vs. undifferentiated (+W/R) states in each stomach region with a less stringent threshold. Delta beta is plotted against the negative logarithm of the raw p-values. The dashed horizontal and vertical lines indicate cutoffs of $p < 0.001$ and delta beta between -0.05 and 0.05, respectively. Red dots refer to hypermethylated CpGs and blue dots to hypomethylated CpGs. **D)** Venn diagrams showing overlaps of DM CpGs upon differentiation in the antrum, corpus, and fundus. The tables provide further details on the overlapping DM CpGs, including stem cell-specific genes (*TNFRSF19* = Troy).

#### Supplementary Figure 2

**A)** Table of the numbers of hypermethylated and hypomethylated genes and promoters, comparing regions within the stomach and the stomach to adjacent tissues. Bottom: Venn diagrams showing overlaps of DM genes and promoters between the stomach regions. **B)** Differential gene expression of DM genes between the stomach regions. Shown are genes with log2 fold changes > 0.6 or < - 0.6, * $p < 0.01$, ** FDR < 0.05. **C)** DM affects genomic features: comparisons of mucosoid samples from different stomach regions (left) and between the stomach to adjacent tissues (right). The plots show the Pearson residuals of the chi-square statistic of counts of significant DM CpGs  located in in TSS1500, TSS200, 5'UTR, 1stExon, gene body, 3'UTR, and IGRs with respect to CpGs islands, -shores, -shelves, and opensea. The dashed line marks the threshold of 95 % confidence interval. TSS1500 and TSS200: 200-1500, and 0-200 bases upstream of the transcription start site (TSS); 5'UTR - 5' untranslated region; Body - gene body; 1stExon - first exon; 3'UTR - 3' untranslated region; IGR - intergenomic region.

#### Supplementary Figure 3

Differential expression of DM TFs between the antrum and the corpus, based on several *in vitro* and *in vivo* data sets (Table 3). The expression level is relative to the antrum mean; the p-values of a student t-test between antrum and corpus are shown on each plot, except for cases with less than two samples in each group. For each study, all micro array probes/probe sets mapping to the respective gene are shown.

#### Supplementary Figure 4

**A)** Gene set enrichment analysis of genes overexpressed in antrum (left) or corpus (right) biopsies compared to differential gene expression between antrum and corpus mucosoids . +W/R and -W/R samples of each region were combined for the mucosoid analysis. Moderated t-scores were used for differential expression ranking. Gene sets of biopsies were determined from (Nookaew et al., 2013, Table 3). ES - enrichment score; NES - normalized enrichment score. **B)** Boxplots of biopsy (red) and mucosoid (green) log2 gene expression fold changes, comparing the corpus and the antrum of selected cell type-specific genes. The dashed line marks the threshold of log 2 fold change > 0.6 or < -0.6. Biopsy samples were taken from Nookaew et al., 2013 (Table 3). **C)** Venn diagrams showing the overlaps of significant upregulated (top) and downregulated (bottom) genes, comparing -W/R (differentiated) to +W/R (undifferentiated) in the different stomach regions (FDR < 0.2; log2 fold change > 0.6 or < -0.6). Numbers in brackets indicate the number of genes belonging to the GO Wnt signaling pathway gene set, and the corresponding genes are listed accordingly. **D)** Normalized expression boxplots of Wnt signaling pathway genes. The boxplots display the median, minimum and maximum normalized expression values of three biological replicates. **E)** Heatmap of *TCF7L1* DNA methylation. Displayed are the normalized methylation values (beta) ranging from 0 (blue, unmethylated) to 1 (red, methylated). Significantly DM CpGs between the antrum and the fundus are marked with red arrows. The black arrow refers to the transcriptional direction. CpG island (CGI).

#### Supplementary Figure 5

**A)** Gene expression log2 fold changes of the stomach- and intestine-specific genes comparing IM and normal organoids (FDR < 0.05). The dashed line indicates a cutoff of log2 fold change > 0.6 or < -0.6. **B)** DNA methylation MDS plot of healthy mucosoids from the antrum, corpus, and fundus, or normal, atrophic, and intestinal-metaplasia (IM) organoids. **C)** Volcano plot of IM vs. normal organoids. Changes in methylation (delta beta) are plotted against the negative logarithm of p-values. The red horizontal and the dashed vertical lines indicate cutoffs of p < 0.05 and delta beta between -0.2 and 0.2, respectively. **D)** Enrichment of sets of CpGs hypermethylated (left) or hypomethylated (right) between IM biopsies and normal biopsies compared to differentially methylation of CpGs between IM and normal organoids. Moderated t-scores were used for ranking the CpGs. Hyper, Hypo indicate the direction of differential methalytion in the IM vs. normal organoid comparison. **E)** Number of DM CpGs, genes, promoters, and enhancers determined in the comparison of IM samples to healthy samples. Left: organoid and biopsy antral IM samples, right: the corresponding samples of the high methylation clusters. **F)** Dot plots of methylation levels (beta values) of the genes *TRIM15* and CDX2: healthy organoid and biopsy samples (top) and IM organoid and biopsy samples (bottom). Squares and crosses denote organoid and biopsy data, respectively. Colors below the plot show DM calls of IM vs. healthy in both data sets.

#### Supplementary Figure 6

**A)** Comparison of organoid IM samples to normal organoid samples - enrichment or depletion of hypermethylated and hypomethylated CpGs in various genomic features. Genomic features were defined based on Encode segmentation classes and transcription factor binding sites (Siggens and Ekwall, 2014), and Roadmap histone modification in gastric tissues (Roadmap Epigenomics Consortium et al., 2015). Shown are log odds ratios of significantly (FDR < 0.05) enriched (log odds ratio > 0.6) or depleted (log odds ratio < -0.6) DM CpGs. The color code ranges from blue (depleted) to red (enriched), grey refers to a log odds ratio < 0.6 or > -0.6. Pol2 - Polymerase 2 subunit; Hyper - hypermethylated CpGs; Hypo - hypomethylated CpGs. **B)** Boxplots of ExE CpG island (CGI) hypermethylation. Shown are the mean ExE CGI hypermethylation values per sample. Healthy samples include, in addition to those of the respective data sets, Roadmap epigenome healthy samples (n = 77), except for the extraembryonic trophoblast cell line and placenta samples. Diseased samples include the respective mild IM and IM (biopsies), IM (organoids), and BE and GC (both biopsies). P values displayed refer to the Wilcoxon rank-sum test with continuity correction. **C)** Histograms of the methylation distribution in ExE hypermethylated CGI and all other CGI taken from the healthy mucosoid data set of the stomach regions and the data set of normal and IM samples from biopsies. The red line and number indicate the median methylation value. **D)** UCSC Genome Browser Image of the *MLH1* promoter region with custom tracks showing the hypermethylation sites in samples of *ex vivo* IM and MSI type of GC (Kent et al., 2002). The sites responsible for *MLH1* silencing in cancer (Morak et al., 2008) are marked in red. Additional tracks show UCSC CpG islands (green) and the UCSC layered H3K27Ac marks.
