## Additional Table 4 for "DNA methylation in human gastric epithelial cells allows cell type-related plasticity and defines regional identity"

| **Gene** | **Site (hg19 coordinates)** | **DM change** | **TF Binding sites** | **Type** | **Enhancer GeneHancer ID** | **Validation, in vivo** | |
| --- | --- | --- | --- | --- | --- | --- | --- |
|  |  |  |  |  |  | **Huang et al.** | **Healthy A/C*** |
| SIM2 | chr21:38119206-38120901 | C > A | EZH2,SUZ12 | 3’ Enh | SIM2/GH21J036746 | Yes | Yes |
| MEIS1 | chr2:66659536-  66659590 | C/F < A | EZH2 | Prom | GH02J066430 | Yes | Yes |
|  | chr2:66810487-66810606 | C/F < A | EZH2 | 3’ Enh | GH02J066572 | Yes | Yes |
| MEIS2 | chr15:37172509-37173017 | C/F < A | EZH2, CDH1 | 3’ CpG island |  | Yes | Yes |
|  | chr15:37180889-37181529 | C/F > A | EZH2, PBX3 | 3’ end |  | Yes | Yes |
|  | chr15:37384042-37389585 | C/F > A | EZH2, others | Prom/Enh | GH15J037092 | Yes | Yes |
|  | chr15:37395115 | C/F > A | EZH2 | Prom/Enh | GH15J037092 | Yes (larger region until 37399798) | Yes (larger region until 37399798) |
| CDX2 | chr13:28541142-28541477 | C < A | EZH2 | Prom | GH13J027958 | Partially | Yes |
|  | chr13:28544760-28547617 | C < A | EZH2,SUZ12 | Prom | GH13J027958 | Yes | Yes |
|  | chr13:28551145 | C < A | EZH2, SUZ12 | Prom | GH13J027958 | Yes | Yes |
| PDX1 | chr13:28492669 | C > A | EZH2,SUZ12 | 5’ Enh | GH13J027917 | Yes | Yes |
|  | chr13:28498956-28503508 | C < A | EZH2, SUZ12 | 3’ end |  | Yes | Yes |
| IRX2 | chr5:2739131 | C > A | EZH2,SUZ12 | 3’ Enh | GH05J002738 | No | Yes |
|  | chr5:2741047 | C > A |  | 3’ Enh | GH05J002738 | Yes | Yes |
|  | chr5:2740667 | C < A | EZH2, CDH1 | 3’ Enh | GH05J002738 | Yes | Yes |
|  | chr5: 2749302 | C < A | EZH2 | Prom | GH05J002746 | No | Yes |
| IRX5 | chr16:54966471-54973385 | C > A | EZH2,SUZ12 | Prom | GH16J054926 | Partially | Partially |
| GATA4 | chr8:11536551 | C > A | EZH2 | 1^st^ Intron |  | No (Cardia only) | Yes |
|  | chr8:11537430-11539405 | C > A | EZH2,SUZ12 | Enh | GH08J011679 | No (Cardia only) | Yes |
|  | chr8:11557597-11560851 | C > A | EZH2,SUZ12 | Prom | GH08J001696 | No (Cardia only) | Yes |
|  | chr8:11565277-11568356 | C > A | EZH2,SUZ12 | Prom | GH08J001696 | No (Cardia only) | No |

Healthy A/C: Methylation data of human stomach antrum biopsies from Yamashita et al. 2019, corpus biopsies from Woo et al. 2018
