## Additional Table 7 for "DNA methylation in human gastric epithelial cells allows cell type-related plasticity and defines regional identity"

**Supplemental Table 7. Data sets created and used in this manuscript**

| Name of the data set | Content | description | GEO accession/  reference |
| --- | --- | --- | --- |
| This study (Mucosoid/ in vitro stomach samples) | DNA methylation and gene expression of healthy gastric epithelial cells | 18 samples  3 biological replicates of   - antrum +W/R and ‑W/R - corpus +W/R and ‑W/R - fundus +W/R and ‑W/R | GSE141660 |
| Healthy stomach in vivo | DNA methylation | 61 normal biopsies   - 39 antrum - 11 corpus - 11 cardia | A subset of GSE103186;  (Huang et al., 2018)^111^[1][1](Huang et al. 2018)(Huang et al. 2018)(Huang et al. 2018) |
|  | Gene expression | 6 normal biopsies (*H. pylori* negative)   - 3 antrum - 3 corpus | A subset of GSE27411 (Nookaew et al., 2013) |
|  |  | 15 normal biopsies   - 11 antrum - 4 corpus | A subset of GSE78523 (Companioni et al., 2017) |
| Healthy stomach in vitro | Gene expression | 3 samples of isolated glands   - 1 antrum - 2 corpus   4 samples of organoids   - 2 antrum - 2 corpus | A subset of GSE60557 (Bartfeld et al., 2015) |
| Combined healthy tissue data set | DNA methylation | 11 normal esophagus biopsies (non-cancer patients) | A subset of GSE72872;  (Krause et al., 2016)^222^[2][2](Krause et al. 2016)(Krause et al. 2016)(Krause et al. 2016) |
|  |  | 2 technical replicates of normal small intestine biopsies | GSE67484;  Unpublished; (Habano 2015) |
|  |  | 3 samples of normal colon biopsies | GSE81211;  (Krause et al., 2016) |
|  |  | 18 mucosoid samples (see above) | GSE141660 |
| This study (normal ex vivo, atrophy, and IM) | DNA methylation and gene expression | 6 samples  2 biological replicates of   - Normal - Atrophic - IM | GSE141660 (DNA methylation and gene expression) |
| Intestinal metaplasia (IM) in vivo | DNA methylation | 191 samples   - 39 normal antrum - 76 IM antrum - 11 normal corpus - 11 normal cardia - 22 mild IM antrum - 23 IM corpus - 9 IM cardia | A subset of GSE103186;  (Huang et al., 2018) |
| Antrum in vivo | DNA methylation | 8 samples of antral biopsy (*H. pylori* negative) | A subset of GSE92863 (Yamashita et al., 2019) |
| Corpus in vivo | DNA methylation | 14 samples of corpus biopsy (*H. pylori* negative) | A subset of GSE99553 (Woo et al., 2018) |
| Barretts’s esophagus (BE) | DNA methylation | 83 samples   - 64 normal esophagus (BE- and cancer patients) - 11 normal esophagus (non-cancer patients) - 19 BE | A subset of GSE72872;  (Krause et al., 2016) |
| Combined healthy samples *in vitro*, IM *ex vivo*, and TCGA STAD data set | DNA methylation | 244 samples   - 18 mucosoid samples - 6 ex vivo samples - 220 TCGA GC samples   - 45 MSI (22 antrum, 20 corpus, 3 GECA)   - 48 GS (24 antrum, 17 corpus, 7 GECA)   - 105 CIN (38 antrum, 44 corpus, 23 GECA) | A subset of the TCGA STAD data set (excluding EBV)(Cancer Genome Atlas Research Network, 2014) |
| GTEx | Gene expression of STAD | 139 normal stomach samples in comparison with TCGA STAD | (GTEx Consortium et al., 2017) |
| Human Protein Atlas | Stomach specific genes | 166 genes with five-fold higher mRNA levels compared to averaged levels in all other tissues | (Uhlén et al., 2015) |
| Chief and parietal GE | Chief and parietal cell signature | Mice chief and parietal cells obtained from laser Capture Microdissection | A subset of GSE5018 (Ramsey et al., 2007) |
| Troy^+^ cells GE | Troy+ cell signature | Significantly enriched genes in Troy+ chief cells compared to whole corpus gland (mice) | Supplementary Table1 (Stange et al., 2013) |
| Beta Catenin target genes | Beta Catenin target genes | β-catenin target genes in human colon or colon carcinoma cells (including colon carcinoma cell lines) | Table 1 (Herbst et al., 2014) |

W/R = Wnt and R-spondin; IM = Intestinal metaplasia; BE = Barrett’s esophagus; STAD = stomach adenocarcinoma.

**Table References:**

1.	Huang KK, Ramnarayanan K, Zhu F, et al. Genomic and Epigenomic Profiling of High-Risk Intestinal Metaplasia Reveals Molecular Determinants of Progression to Gastric Cancer. Cancer Cell 2018;33:137-150 e5.

2.	Krause L, Nones K, Loffler KA, et al. Identification of the CIMP-like subtype and aberrant methylation of members of the chromosomal segregation and spindle assembly pathways in esophageal adenocarcinoma. Carcinogenesis 2016;37:356-65.
